# Insecticides can simultaneously target mosquito vectors and malaria parasites

**DOI:** 10.64898/2026.06.15.732335

**Authors:** Antonia L Böhmert, Moritz Sturm, Natalie M Portwood, Julia B Mäurer, Friedrich Frischknecht, Fred Hamprecht, Victoria A Ingham

## Abstract

Insecticide-based vector control remains the cornerstone of malaria prevention, averting approximately 1.2 billion cases between 2000 and 2025. These interventions primarily reduce transmission by killing mosquitoes; however, widespread reliance on a limited number of compounds has driven the emergence of insecticide resistance. This has prompted the development of new insecticides with novel modes of action. Notably, the pyrrole insecticide chlorfenapyr has been shown to affect both the mosquito vector and the malaria parasite, suggesting that compounds with dual activity could provide an additional strategy to suppress transmission. Here, we present a medium-throughput discovery pipeline that integrates *in vitro Plasmodium* sporozoite motility assays with machine-learning-based analysis, alongside *in vivo* exposure of infected *Anopheles* mosquitoes and quantification of parasite development. Screening 32 insecticidal chemistries identified five compounds that significantly impaired sporozoite motility, including three avermectin endectocides, the mitochondrial complex III inhibitor hydramethylnon, and tralopyril, the active form of chlorfenapyr. Several compounds transiently increased motility, indicating that parasite physiology is frequently influenced by insecticide exposure. *In vivo* exposure to abamectin reduced parasite numbers in both the haemolymph and salivary glands and impaired productive motility. Importantly, this inhibition was confirmed in *Plasmodium falciparum*-infected mosquitoes, where exposure significantly reduced salivary gland invasion.

These findings reveal that parasite-directed activity among insecticides may be more common than previously appreciated and demonstrate a scalable approach to identify compounds capable of simultaneously killing mosquitoes and suppressing parasite transmission.

**Significance Statement:** Vector control relies heavily on insecticides that kill mosquitoes, yet rising resistance threatens their effectiveness. Here we show that several insecticides also affect the malaria parasite itself. Using a scalable screening pipeline combining machine learning–assisted sporozoite motility analysis with mosquito infection assays, we found that 15% of tested insecticides significantly impaired parasite motility, including compounds with distinct modes of action. Among these hits, the avermectin abamectin reduced parasite dissemination in mosquitoes and limited salivary gland invasion in both *Plasmodium berghei* and the human malaria parasite *P. falciparum*. These findings reveal that parasite-directed activity among insecticides may be more widespread than expected and highlight the potential to develop vector control tools that simultaneously kill mosquitoes and block parasite transmission.

## Introduction

Malaria remains a significant global health threat, with an estimated 282 million cases and 610,000 deaths in 2024 alone (WHO, 2025b). Malaria is a vector-borne disease caused by five species of the apicomplexan parasite *Plasmodium* of which *P. falciparum* is responsible for the majority of severe disease and mortality. Transmission occurs through the bite of infected *Anopheles* mosquitoes and is primarily controlled through insecticide-based interventions such as insecticide treated bed nets (ITNs) and indoor residual spraying (IRS). Together, ITNs and IRS averted over 1.2 billion cases since the year 2000, corresponding to 75% of averted cases (Malaria Atlas Project, 2025), making them by far the most effective malaria control intervention. However, the limited number of chemical classes available, particularly for ITNs, the most efficacious tool, has contributed to widespread insecticide resistance in *Anopheles* populations (Ranson & Lissenden, 2016; WHO, 2025b), creating an urgent need for new insecticidal chemistries. Bringing novel chemistries to market, however, typically requires close to a decade of research, development, and regulatory evaluation (WHO, 2012). Identifying additional activities within existing or repurposed compounds may therefore provide a faster route to expanding the arsenal of vector control tools (Lees et al., 2019).

Currently, the World Health Organization (WHO) recommends ITNs containing pyrethroids in combination with a second active ingredient (WHO, 2025a). The most important of these combines pyrethroids with the pyrrole insecticide chlorfenapyr, whilst ITNs pairing pyrethroids and the insecticide synergist piperonyl butoxide or the sterilizing agent pyriproxyfen have also received conditional recommendations for use (Maiteki-Sebuguzi et al., 2023; Mosha et al., 2023; Protopopoff et al., 2023; WHO, 2024). IRS allows the use of a broader spectrum of chemistries due to reduced human contact including pyrethroids, carbamates, organophosphates, neonicotinoids and recently the meta-diamide broflanilide, the isooxazoline isocycloseram and chlorfenapyr (Blythe et al., 2022; Ngufor et al., 2021; WHO, 2024). Following a blood meal, female mosquitoes resting on a treated surface contact these insecticides, which induces lethal responses in the mosquito, thereby preventing potential transmission of the malaria parasite from one host to the next. Most currently deployed insecticides are neurotoxic, targeting voltage-gated ion channels (pyrethroids) (Narahashi, 1971), acetylcholine esterase (organophosphates and carbamates) (Kuhr, 1976; Mileson et al., 1998), the nicotinic acetylcholine receptor (neonicotinoids) (Taillebois, Cartereau, Jones, & Thany, 2018), the ryanodine receptor (diamides) (Du & Fu, 2023), or GABA-gated chloride channels (isoxazolines, meta-diamides) (Blythe et al., 2022; Zeng et al., 2026), thereby inhibiting synaptic transmission or causing overstimulation, leading to temporary knockdown and eventual death (Ndiath, 2019). In contrast, chlorfenapyr acts through a distinct mechanism by disrupting mitochondrial oxidative phosphorylation (Black, 1994). As this pathway is conserved across eukaryotes, such modes of action raise the possibility that insecticides may directly influence parasite development within the mosquito.

Within the mosquito vector, *Plasmodium* undergoes several developmental transitions that represent critical transmission bottlenecks. Following ingestion of typically thousands of gametocytes during a blood meal, parasites undergo fertilisation in the mosquito midgut, forming <100 motile ookinetes which traverse the gut epithelium and develop into oocysts, often no more than 10 per mosquito (Gouagna et al., 1998; Smith, Vega-Rodriguez, & Jacobs-Lorena, 2014). Subsequent sporogony produces thousands of sporozoites that egress into the haemolymph, of which a small fraction successfully invades the salivary glands, from where they are transmitted to a new host during the next blood meal (Kanatani, Stiffler, Bousema, Yenokyan, & Sinnis, 2024; Sinden & Croll, 1975; Smith et al., 2014; Venugopal, Hentzschel, Valkiunas, & Marti, 2020). Each of these stages is characterised by rapid cellular differentiation, intense metabolic activity, and exposure to mosquito immune responses, making parasite development within the mosquito susceptible to environmental perturbations.

Recent evidence suggests that insecticide exposure can influence parasite development and transmission within the mosquito. For example, pyrethroid exposure has been shown to increase reactive nitrogen species responses in *Anopheles* mosquitoes, leading to enhanced ookinete lysis and reduced infection prevalence (Hörner, 2025), demonstrating that even neurotoxic insecticides may indirectly affect parasite survival. Conversely, the pyrrole insecticide chlorfenapyr has been reported to directly impair parasite development through its active metabolite tralopyril, which disrupts mitochondrial respiration and blocks oocyst formation in both *P. berghei* and *P. falciparum* (Kweyamba et al., 2023; Portwood, 2026). These findings are consistent with the well-established susceptibility of mosquito-stage parasites to mitochondrial inhibition, as illustrated by antimalarial compounds such as atovaquone that target the parasite electron transport chain and can prevent parasite transmission (Painter, Morrisey, & Vaidya, 2010; Paton et al., 2019; Probst et al., 2025; Srivastava, Rottenberg, & Vaidya, 1997). Together, these studies suggest that insecticides may influence malaria transmission not only by killing mosquitoes but also by directly or indirectly affecting parasite development. Exploiting such dual activity could provide a powerful strategy for vector control, particularly in the context of widespread insecticide resistance. However, systematic approaches to identify compounds with parasite-directed activity remain limited.

Here we reveal that insecticides can frequently exert dual activity against both the mosquito vector and the malaria parasite. Screening insecticidal chemistries identified a surprising proportion that modulated *Plasmodium* sporozoite motility, including the avermectins abamectin, ivermectin, and emamectin benzoate, the mitochondrial complex III inhibitor hydramethylnon, and tralopyril, the active metabolite of the pyrrole insecticide chlorfenapyr, which inhibited motility. These effects were observed in the rodent malaria parasite *P. berghei* and confirmed for abamectin in the predominant human malaria parasite *P. falciparum*. By integrating machine-learning-assisted sporozoite motility analysis with *in vivo* mosquito infection assays, our medium-throughput screening pipeline enables rapid identification of vector control compounds with previously unrecognised parasite-directed activity. These findings suggest that parasite susceptibility to insecticides may be far more common than previously appreciated and highlight the potential to develop vector control chemistries that simultaneously kill mosquitoes and suppress parasite transmission.

## Results

### Automated sporozoite motility classification enables medium-throughput screening of dual activity

Mosquitoes may encounter insecticides whilst carrying mature salivary gland sporozoites from a prior infection, thus permitting a direct interaction of the sporozoites with the compounds or an indirect effect through insecticide-induced physiological change. To systematically address the former, insecticidal compounds were tested for their effect on sporozoite motility (Figure 1A), a key determinant of infection (Amino et al., 2006; Hopp et al., 2015), which is driven by an actomyosin motor system that mediates the rearward translocation of surface adhesins such as thrombospondin-related anonymous protein (TRAP), enabling forward movement (Kappe, Buscaglia, Bergman, Coppens, & Nussenzweig, 2004; Meissner, Ferguson, & Frischknecht, 2013). We employed a modified version of a previously described assay (Prinz, Sattler, & Frischknecht, 2017), exposing sporozoites to insecticides at physiologically relevant concentrations (Spielmeyer, Schetelig, & Etang, 2019) (Supplementary Table 1). Specifically, we focused on perturbations of circular, productive motility and the induction of various non-productive motility types described previously (Prinz et al., 2017) (Figure 1B).

**Figure 1.**
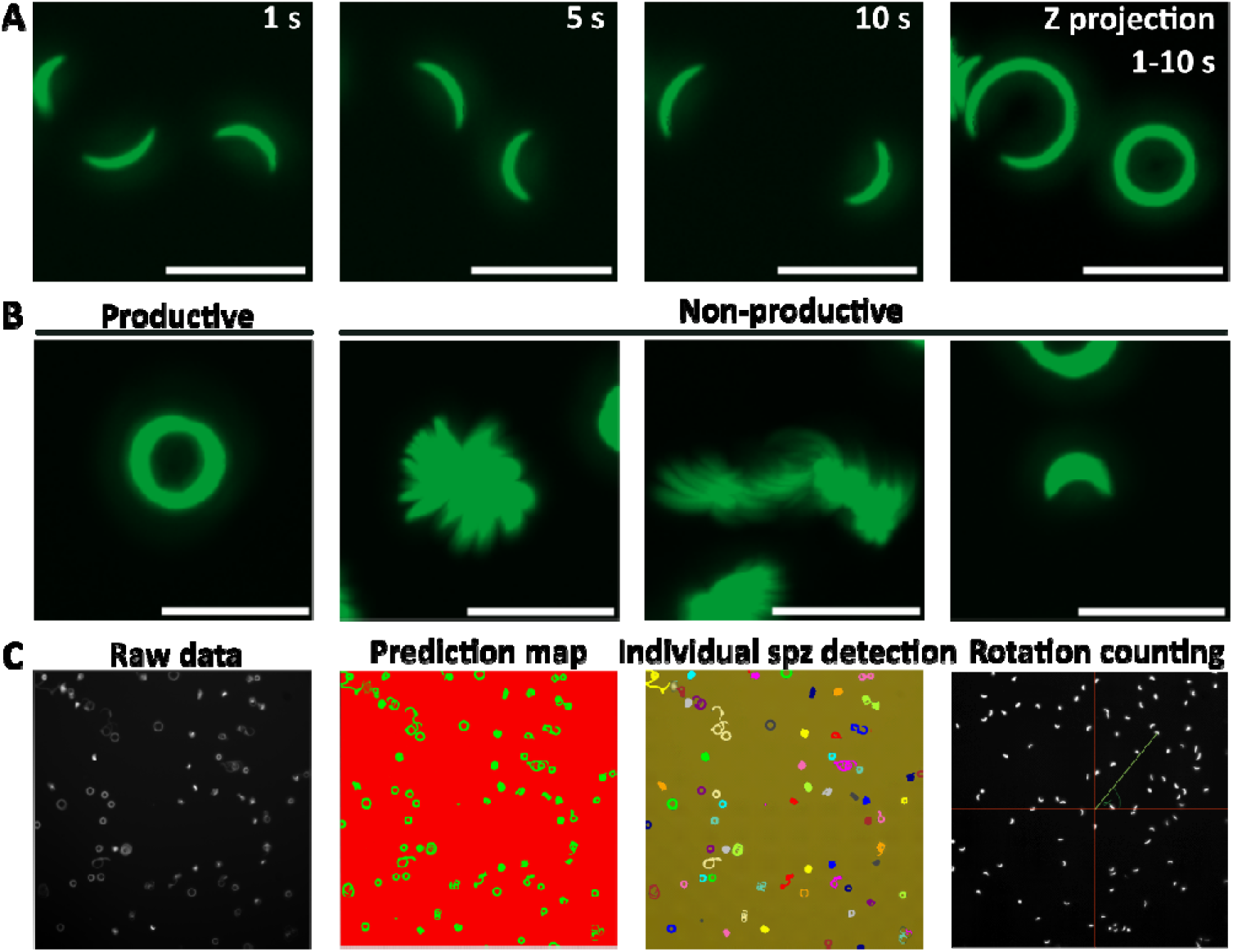
*Plasmodium* 2D sporozoite motility *in vitro*. (A) GFP-positive *P. berghei* sporozoites migrate on a flat glass-bottom surface in a counter-clockwise orientation, with their circular trajectories elucidates by a maximum intensity z projection of timepoints 1 s, 5 s, and 10 s. Scale bar = 20 µm. (B) Representative images of maximum intensity z projection of sporozoites undergoing productive motility (left), which we consider productive, vs. various non-productive motility patterns (right). Scale bar = 20 µm. (C) Workflow of *Ilastik*-based sporozoite tracking and labelling, as well as rotation counting of productively migrating sporozoites (Sommer C., 2011). Using the raw data, a prediction map of sporozoites (green) vs. background (red) is generated, and individual sporozoites detected (multiple colours) and assigned a motility pattern based on training with a subset of motility recordings. Finally, the number of rotations of sporozoites labelled as moving in a circular fashion was assessed based on oscillations in angle of sporozoites compared to a fixed position in the field of view.

To increase analytical throughput and minimise user bias during manual labelling, we developed an automated motility classification pipeline using *Ilastik*-based machine learning (Sommer C., 2011). The classifier was trained on a curated subset of sporozoite motility recordings generated in this study and enabled automated assignment of motility phenotypes (Figure 1C). Inter-user variability between two trained annotators ranged from 1–8%, while the variability between a trained user and the tool was 5-15% (Supplementary Figure 1). Automating motility classification substantially increased analytical efficiency and enabled standardised quantification of productive motility across compound treatments.

### *In vitro* insecticide exposure can impact sporozoite motility

Using an adapted version of a previously described motility assay (Prinz et al., 2017) (Figure 2A), 32 compounds belonging to 15 Insecticide Resistance Action Committee (IRAC) groups, including active insecticides and pro-insecticides, as well as active metabolites, synergists, herbicides and unlicensed compounds, were screened (Figure 2B, Supplementary Table 1, Supplementary Table 2). Whilst many of the tested compounds did not significantly impact sporozoite motility, we observed a significant decrease in productive motility upon exposure to five out of 32 compounds (15%). These included all tested members of the avermectin endectocides: abamectin, ivermectin and emamectin benzoate, as well as tralopyril, the active form of the pyrrole chlorfenapyr (Black, 1994), as previously demonstrated (Portwood, 2026), and the mitochondrial complex III electron transport inhibitor hydramethylnon. Exposure to hydramethylnon induced both a time- and dose-dependent effect, with a significant reduction in productive motility observed 10 min post-exposure at 13.5 µg/ml (*p*_2-way ANOVA_ = 0.0027), whereas by 60 min all tested concentrations significantly impaired motility (0 min *p*_2-way ANOVA_ = 0.0078, 20 min *p*_2-way ANOVA_ = 0.0005, 60 min *p*_2-way ANOVA_ = 0.0003). Tralopyril similarly induced a time-dependent effect, with significant reduction in motility at all tested concentration from 10 min onwards (9 µg/ml *p*_2-way ANOVA_ = 0.0032). Interestingly, ivermectin and abamectin caused an immediate and nearly complete loss of productive motility at 9 µg/ml (abamectin *p*_2-way ANOVA_ = <0.0001; ivermectin *p*_2-way ANOVA_ = <0.0001). Emamectin-benzoate also acted immediately but displayed a clear dose-dependence, with significant inhibition only at 22.5 µg/ml (*p*_2-way ANOVA_ = <0.0001). Two additional compounds, the acetyl-CoA carboxylase inhibitor clodinafop propargyl and the mitochondrial complex II inhibitor cyflumetofen, showed trends towards reduced motility relative to the untreated controls (clodinafop propargyl, 9 µg/ml, 60 min *p*_2-way ANOVA_ = 0.275; cyflumetofen, 13.5 µg/ml, 60 min *p*_2-way ANOVA_ = 0.0571).

**Figure 2.**
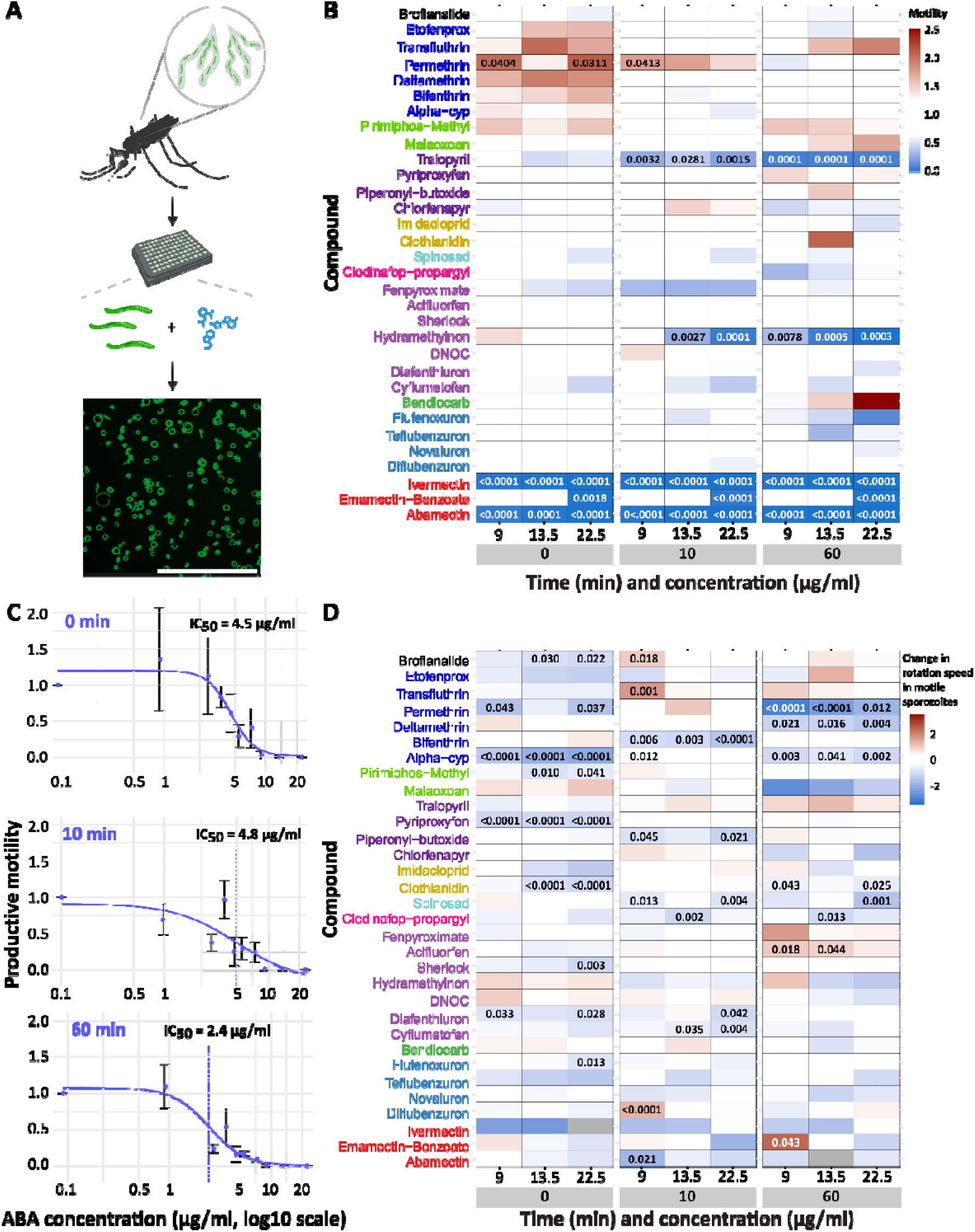
Insecticides impact *P. berghei* sporozoite motility *in vitro*. **(A)** Schematic of *in vitro* insecticide exposure of csGFP *P. berghei* sporozoites for effects on sporozoite motility. Scale bar = 300 µm. **(B)** *In vitro P. berghei* sporozoite motility in response to *in vitro* exposure to compounds. The ratio of productive motility compared to the respective control is represented from reduced (<1, blue) to increased (>1, red) for timepoint 0-, 10- and 60-min post-addition of the compound at 9, 13.5 and 22.5 µg/ml. Compound names are coloured by mode of action: GABA modulators (black), pyrethroids (blue), organophosphates (green), new-net compounds (dark purple), neonicotinoids (yellow), benzoylureas (light blue), Acetyl-CoA-carboxylase inhibitors (pink), mitochondrial inhibitors (purple), carbamates (dark green), inhibitors of chitin biosynthesis (cyan), and avermectins (red). n= 32 compounds tested across 3 concentrations and 3 timepoints. Three biological replicates of n= 100-200 sporozoites per sample were run for each compound. Significant differences in productive motility compared to the control were determined by two-way ANOVA with multiple comparisons, with significant p-values displayed within cells. **(C)** Dose response of *P. berghei* sporozoite productive motility following *in vitro* abamectin (ABA) exposure at 0-, 10-, and 60-min post-addition. Data points represent means and error bars standard deviation. IC_50_ values at each timepoint are indicated by the dashed line. Three biological replicates of n= 100-200 sporozoites per sample were run for each compound. Significant differences in productive motility compared to the control were determined using a Welch’s t-test, with concentrations with significant p-values (<0.05) shaded in grey. **(D)** Effects of insecticide exposure on sporozoite rotation speed among productively motile sporozoites. Linear mixed-effects modelling was used to assess gliding speed differences between treated and control conditions, restricted to sporozoites exhibiting productive motility. Heatmap shows model estimates representing the difference in mean number of migrated circles per sporozoite between each treatment condition and its corresponding control (0 µg/ml) at the same timepoint. Blue indicates reduced migration speed relative to control; red indicates increased speed. Grey cells indicate insufficient productive sporozoites for statistical analysis. Significant differences after Benjamini-Hochberg false discovery rate correction (FDR < 0.05) are indicated by p-values displayed within cells. Compound names are colour-coded as in (B). Data represent pooled analysis of individual sporozoite rotation counts from three independent biological replicates. n = 32 compounds tested across 3 concentrations and 3 timepoints.

Given the clear dose dependence observed for the avermectins, we next determined the IC_50_ for inhibition of productive motility. Abamectin displayed an exceptionally potent phenotype, inducing complete loss of motility immediately upon exposure at concentrations ≤4.5 µg/ml (*p*_2-way ANOVA_ = 0.0016) (Figure 2C, Supplementary Table 3). After 60 min, the IC_50_ was 2.4 µg/ml, underscoring the high intrinsic susceptibility of sporozoites to this compound and suggesting that even low dose sub-lethal exposure within the mosquito could substantially impair parasite transmission. Indeed, as the assay duration is limited to 60 min, beyond which activated sporozoites progressively lose motility independent of treatment (Hopp et al., 2015), this estimate likely represents a conservative measure of potency.

Interestingly, pyrethroid insecticides generally induced a transient increase in productive sporozoite motility immediately after exposure; this effect was short-lived and largely disappeared within 10 min (Figure 2B). Permethrin was the notable exception, producing a significant increase in productive motility at the lowest concentration at 10 min (*p*_2-way ANOVA_ = 0.0413) and a broadly increased response immediately upon exposure (9 µg/ml *p*_2-way ANOVA_ = 0.0404; 22.5 µg/ml *p*_2-way ANOVA_ = 0.0311).

The pipeline also quantified the number of migrated parasite circles as a proxy for sporozoite speed. Of the 288 possible comparisons (32 compounds × 9 conditions), 52 showed significant differences from control after FDR correction (Figure 2D, Supplementary Table 4). Alpha-cypermethrin demonstrated the most pronounced and consistent effects, with significant reductions in speed observed at all timepoints and concentrations. Permethrin showed particularly strong effects at later timepoints, with the most pronounced reductions at 60 min. Clothianidin exhibited concentration-dependent effects primarily at early exposure, with significant reductions at 0 min and 10 min at ≥ 13.5 µg/ml and 22.5 µg/ml, respectively. Several pyrethroids showed time-dependent responses, with bifenthrin effects most apparent at 10 min at ≥9 µg/ml, while deltamethrin, permethrin and alpha-cypermethrin showed delayed effects emerging primarily at 60 min at ≥9 µg/ml. Spinosad demonstrated concentration-dependent inhibition at 10 min and strong effects at the highest concentration by 60 min. Abamectin induced a significant decrease at 10 min at 9 µg/ml, while emamectin benzoate induced a significant increase only at 60 min at 9 µg/ml. Eleven compounds showed no significant effects on rotation speed among productive movers after FDR correction.

### *In vivo* abamectin but not hydramethylnon exposure of *P. berghei*-infected mosquitoes reduces sporozoite egress from oocysts and salivary gland invasion

To assess the effect of abamectin exposure on maturing parasites, we first established a dose that ensured high insecticide uptake whilst maintaining sufficient mosquito survival. Topical dose-response assays revealed irreversible flaccid paralysis within 24 h post-exposure, a phenotype previously reported for mosquitoes and nematodes (de Freitas, Faria Mde, Alves, & de Melo, 1996; Jeena et al., 2025; Wolstenholme & Rogers, 2005). Although the mosquitoes survived the initial exposure, paralysis prevented sugar feeding, leading to subsequent desiccation and death (Supplementary Figure 2). We therefore selected the LC_30_ at 72 h (5.8 µg/ml) and performed exposures at 10 days post infection (dpi), when oocysts are mature and sporulation has taken place (Dimopoulos, Seeley, Wolf, & Kafatos, 1998; Pimenta, Touray, & Miller, 1994). Parasite quantification was then conducted 3-days post exposure, specifically targeting the window encompassing sporozoite egress from oocysts and salivary gland invasion (Figure 2A).

As expected, oocyst prevalence did not differ between the exposed and control group (*p*_Fisher’s exact test_ = 0.5634) (Figure 2B), with no significant difference in oocyst numbers per midgut (*p*_One-way ANOVA_ = 0.3331) (Figure 3C, Supplementary Figure 3A). However, average oocyst size was significantly increased following abamectin exposure (13% increase, *p*_One-way ANOVA_ = 0.0105) (Figure 3D), a phenotype observed across all individual replicates (Supplementary Figure 3B), indicative of impaired or delayed egress (Portwood, 2026). Indeed, abamectin exposure resulted in significantly fewer burst oocysts (35% reduction, *p*_One-way ANOVA_ = 0.0003) (Figure 3E, Supplementary Figure 3C), and culminated in a non-significant reduction of haemolymph sporozoites (67% reduction, *p*_One-way ANOVA_ = 0.4405) (Figure 3F, Supplementary Figure 3D) and a significant and comparable reduction in salivary gland sporozoite numbers (56% reduction, *p*_One-way ANOVA_ = 0.0005) (Figure 3G, Supplementary Figure 3E). These findings indicate that abamectin disrupts sporozoite egress from the oocyst, thereby limiting subsequent dissemination and gland invasion. In addition, it might also affect salivary gland invasion of egressed sporozoites. Consistent with these quantitative data, the number of fluorescent salivary glands, an indicator of sporozoite presence due to the expression of GFP under the CSP promoter (Natarajan et al., 2001), was significantly reduced in exposed mosquitoes (37% reduction, *p*_Fisher’s exact test_ = <0.0001) (Figure 3H), further supporting diminished sporozoite invasion.

**Figure 3.**
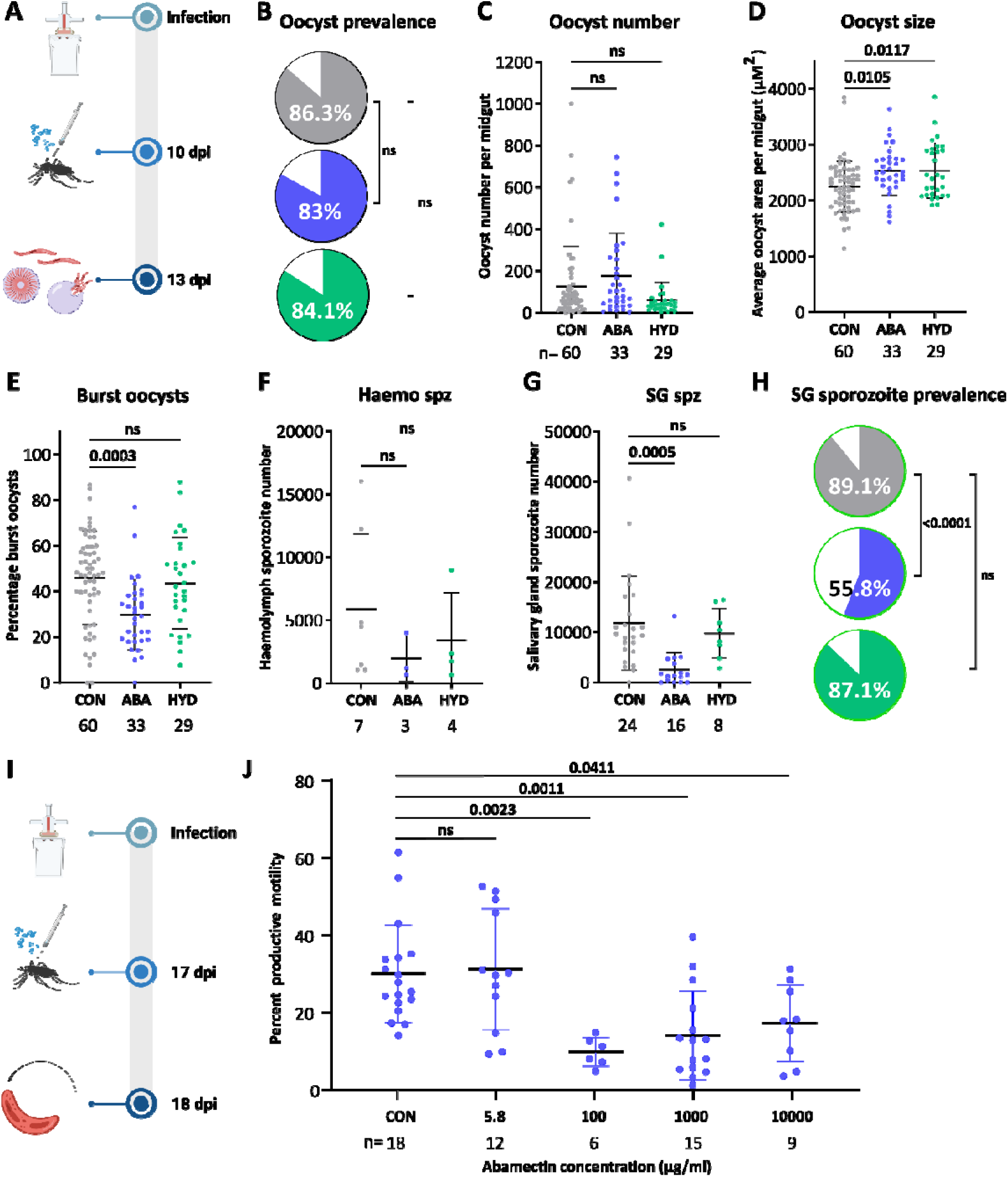
*In vivo* abamectin but not hydramethylnon exposure of *P. berghei*-infected mosquitoes reduces sporozoite egress from oocysts and salivary gland invasion. **(A)** Schematic of *in vivo* exposure of *P. berghei*-infected mosquitoes for oocyst and sporozoite development assessment. Infected mosquitoes were topically exposed 10 days post-infection (dpi) and oocysts and sporozoites assessed three days later at 13 dpi. **(B-H)** *P. berghei* infection responses to abamectin (ABA, blue) and hydramethylnon (HYD, green) vs control (CON, grey) exposure 13 days post-infection and three days post-exposure. Lines represent means and error bars indicate standard deviation. Significant differences in prevalence determined by Fisher’s exact test and for oocyst and sporozoite counts and burst oocysts by one-way ANOVA with multiple comparisons, with p<0.05 considered significant. **(B)** Oocyst prevalence 13 days post-exposure (n CON = 117, ABA = 52, HYD = 63). **(C)** Oocyst number per midgut (n CON= 60, ABA = 33, HYD = 29). **(D)** Mean oocyst size in µm^2^ per midgut (n CON= 60, ABA= 33, HYD = 29). **(E)** Percentage burst oocysts per midgut (n CON= 60, ABA= 33, HYD = 29). **(F)** Mean haemolymph sporozoite number per mosquito (n CON= 7, ABA= 3, HYD = 4; pools of 20 mosquitoes). **(G)** Mean salivary gland sporozoite number per mosquito (n CON = 24, Aba= 16, HYD = 8; pools of 5 salivary glands). **(H)** Salivary gland sporozoite prevalence, based on GFP-signal in salivary glands (n CON = 119, ABA = 43, HYD = 70). **(I)** Schematic of *in vivo* abamectin exposure of *P. berghei*-infected mosquitoes for subsequent *in vitro* sporozoite motility assessment. **(J)** *In vitro P. berghei* sporozoite motility for sporozoites of abamectin-exposed (blue) vs. control-treated (grey) mosquitoes, 24 h post-*in vivo* exposure. Lines represent means and error bars indicate standard deviation. n= 100-200 sporozoites per concentration per replicate (n CON = 18, 5.8 µg/ml = 12 replicates, 100 µg/ml = 6 replicates, 10,000 µg/ml = 15 replicates, 10,000 µg/ml = 9 replicates). Significant differences were determined by two-way ANOVA with multiple comparisons, with p<0.05 considered significant. P-values indicated within figure.

Given its inhibitory effect on sporozoite motility *in vitro*, hydramethylnon was similarly evaluated following *in vivo* exposure with *P. berghei*. After determining the LC_30_ dose at 72 h (200 µg/ml, Supplementary Figure 4), infected mosquitoes were topically exposed 10 days post-infection, mirroring the abamectin experimental design (Figure 3A). While hydramethylnon exposure also impacted oocyst development, sporozoite dissemination was not affected. Oocyst number (*p*_One-way ANOVA_ = 0.1966) (Figure 3C, Supplementary Figure 3A), proportion of burst oocysts (*p*_One-way ANOVA_ = 0.8143) (Figure 3E, Supplementary Figure 3C), as well as haemolymph (*p*_One-way ANOVA_ = 0.6624) (Figure 3F, Supplementary Figure 3D) and salivary gland sporozoite numbers (*p*_One-way ANOVA_ = 0.7210) (Figure 3G, Supplementary Figure 3E) were comparable to controls three days post-exposure, while size was significantly increased (*p*_One-way ANOVA_ = 0.0117) (Figure 3D, Supplementary Figure 3B) putatively indicating that high doses could affect these parameters. Infection prevalence did not differ significantly in the midgut (*p*_Fisher’s exact_ = 0.8433) (Figure 3B) nor in the salivary glands (*p*_Fisher’s exact_ = 0.8282) (Figure 3H). To assess potential earlier effects on zygotes, ookinetes and oocyst formation, exposures were also performed immediately post-infection (Supplementary Figure 5A). No significant differences in prevalence (*p*_Fisher’s exact_ = 0.2828) (Supplementary Figure 5B), oocyst intensity (*p*_Mann-Whitney_ = 0.6060) (Supplementary Figure 5C), or size (*p*_Mann-Whitney_ = 0.2084) (Supplementary Figure 5D) were observed five days post-exposure, indicating no impact on gametocyte activation, ookinete production and traversal or oocyst establishment.

### Abamectin exposure reduces sporozoite motility in *P. berghei-*infected mosquitoes

Next, we assessed the *in vitro* gliding motility of sporozoites 24 h after *in vivo* abamectin exposure to determine whether the *in vitro* data is recapitulated by *in vivo* exposure (Figure 3I). To better assess dose-responsiveness, higher concentrations (100 µg/ml, 1,000 µg/ml, 10,000 µg/ml) were tested alongside the LC_30_ dose (5.8 µg/ml) used in the developmental assays. A consistent and significant dose-dependent reduction in productive motility compared to the untreated control can be observed from >100 µg/ml (100 µg/mL *p*_2-way ANOVA_ = 0.0023; 1,000 µg/ml *p*_2-way ANOVA_ = 0.0011; 10,000 µg/ml *p*_2-way ANOVA_ = 0.0411) (Figure 3J).

### Abamectin exposure reduces *P. falciparum* sporozoite invasion of salivary glands

To test whether abamectin produced comparable transmission blocking efficacy in the human malaria parasite, we repeated the *in vivo* exposure experiments using *P. falciparum* infected mosquitoes. As observed in *P. berghei*, abamectin exposure did not alter infection prevalence (*p*_Fisher’s exact_ = 0.2582) (Figure 4A) or oocyst intensity (*p*_Mann-Whitney_ = 0.3888) (Figure 4B, Supplementary Figure 6A) and oocyst size was unchanged (*p*_Mann-Whitney_ = 0.6442) (Figure 4C, Supplementary Figure 6B), indicating that early sporogonic development was not detectably impaired. Although the proportion of burst oocysts in control mosquitoes was relatively low, suggesting that exposure may have occurred slightly prior to peak egress, no evidence of delayed oocyst maturation was observed (*p*_Mann-Whitney_ = 0.9419) (Figure 4D, Supplementary Figure 6C). Importantly, despite a non-significant reduction in haemolymph sporozoite numbers (44% reduction, *p*_Mann-Whitney_ = 0.2786) (Figure 4E, Supplementary Figure 6D), abamectin exposure resulted in a clear reduction in salivary gland sporozoites (73% reduction, *p*_Mann-Whitney_ = 0.0012) (Figure 4F, Supplementary Figure 6E). Thus, as seen in *P. berghei*, abamectin exposure during late sporogony significantly limits successful salivary gland invasion, directly restricting the transmissible parasite population.

**Figure 4.**
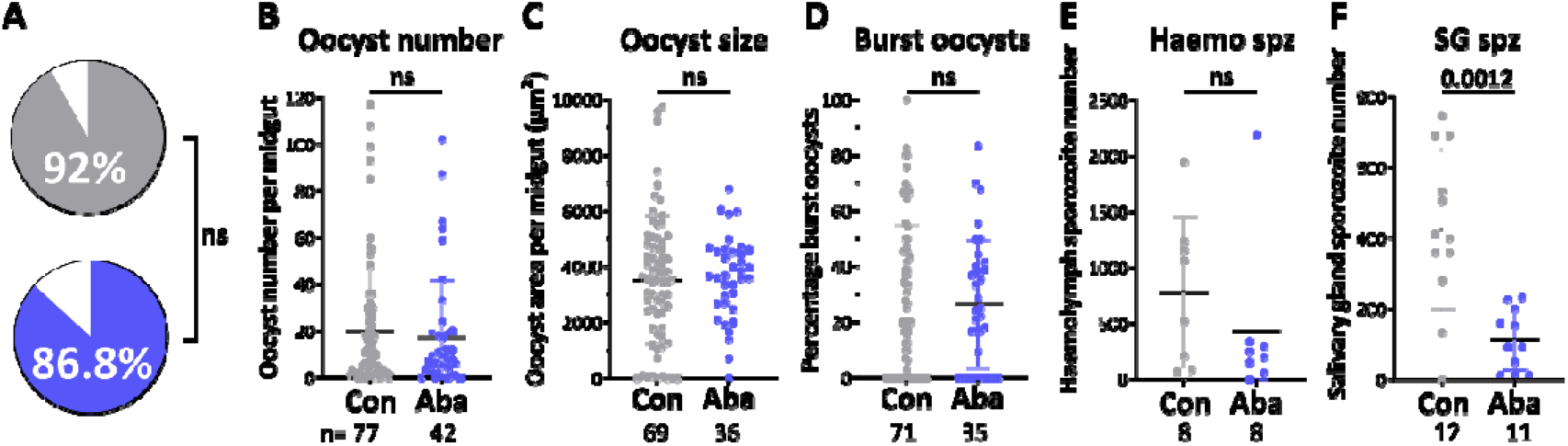
*In vivo* abamectin exposure of *P. falciparum*-infected mosquitoes reduces salivary gland sporozoites. **(A-F)** *P. falciparum* infection responses to abamectin (ABA, blue) vs control (CON, grey) exposure 13 days post-infection and 3 days post-exposure. Significant differences in prevalence determined by Fisher’s exact test and for oocyst and sporozoite counts and burst oocysts by Mann-Whitney test, with p<0.05 considered significant. **(A)** Oocyst prevalence 13 days post-exposure (n CON = 77, ABA = 42). **(B)** Oocyst number per midgut (n CON= 77, ABA = 42). **(C)** Mean oocyst size in µm^2^ per midgut (n CON= 69, ABA= 36). **(D)** Percentage burst oocysts per midgut (n CON= 71, ABA= 35). **(E)** Mean haemolymph sporozoite number per mosquito (n CON= 8, ABA= 8; pools of 2-20 mosquitoes). (F) Mean salivary gland sporozoite number per mosquito (n CON= 12, ABA= 11; pools of 5 salivary glands).

## Discussion

The widespread rise of insecticide resistance in anopheline malaria vector populations, particularly to pyrethroid insecticides, challenges global malaria eradication efforts. Consequently, the identification of novel or repurposed compounds for use in vector control is of great importance. Compounds with a dual effect, targeting both the vector and the parasite, are of particular interest, as they may mitigate the impact of emerging insecticide resistance. Such dual effects may arise indirectly through sub-lethal exposure resulting in physiological changes in the host, which disrupts parasite development, or directly through toxic effects of insecticides or their metabolites on the parasite itself. Here we present a medium-throughput pipeline, integrating *in vitro* sporozoite motility assays with machine-learning-based analysis, alongside *in vivo* exposures of *Plasmodium* infected *Anopheles* mosquitoes. This approach enables the simultaneous assessment of sporozoite motility as a proxy for transmission potential and the evaluation of compound effects on both early and late parasite developmental stages. Using this platform we identified several classes of insecticidal compounds with previously underappreciated transmission-blocking potential.

Among the compounds tested, the avermectins abamectin, ivermectin, and emamectin benzoate emerged as potent dual-activity compounds. These macrocyclic lactones act as modulators of glutamate-gated chloride channels (GluCls), ligand-gated ion channels restricted to protostome phyla, including insects, and absent in vertebrates (Wolstenholme, 2012). By directly activating or potentiating glutamate-dependent channel opening (Cully, Paress, Liu, Schaeffer, & Arena, 1996; Cully et al., 1994) avermectins induce neuronal hyperpolarisation and flaccid paralysis in target organisms. GluCls are well-conserved in nematodes and arthropods (Cully et al., 1994; Glendinning, Buckingham, Sattelle, Wonnacott, & Wolstenholme, 2011; McCavera, Rogers, Yates, Woods, & Wolstenholme, 2009; Wolstenholme & Rogers, 2005), including the malaria vector *An. gambiae* (Meyers et al., 2015), but have not been identified in apicomplexan parasites such as *Plasmodium*, indicating that the anti-plasmodial effects observed here may arise through an alternative mechanism. Previous studies have reported inhibitory activity of ivermectin against human-erythrocytic and hepatic stages of both *P. berghei* and *P. falciparum* (Mendes et al., 2017; Panchal et al., 2014), supporting the concept that avermectins exert parasite-directed effects independent of their insect neuronal targets. One proposed mechanism involves inhibition of importin α/β-mediated nuclear transport, disrupting signal recognition particle (SRP)-associated protein trafficking (Panchal et al., 2014). Avermectins have also been linked to DNA damage and apoptotic death through activation of the oxidative-stress and mitochondrial-stress pathway responses (Zhang et al., 2022). Given the rapid and near-complete loss of productive sporozoite motility observed *in vitro*, interference with essential cellular processes such as protein trafficking or cytoskeletal regulatory pathways is consistent with our data. The pronounced reduction in productive motility was accompanied by only limited effects on migration speed among the few remaining motile sporozoites. While this likely reflects the near-complete loss of motile parasites, which constrains statistical power, it also suggests that avermectins primarily act as a binary switch determining whether sporozoites retain productive motility, rather than modulating the kinematics of movement in migrating parasites. Furthermore, the impaired egress and reduced salivary gland invasion after *in vivo* exposure similarly support a direct impact on parasite biology rather than solely vector-mediated effects.

The active metabolite of the pyrrole insecticide chlorfenapyr, tralopyril, induced a time-dependent reduction in productive sporozoite motility, whereas the pro-insecticide itself had no immediate effect, consistent with the requirement for metabolic activation by cytochrome P450 enzymes within the mosquito vector (Yunta et al., 2023) and absence of characterised P450 enzymes in *Plasmodium* (van Dooren, Kennedy, & McFadden, 2012). Again, sporozoites that retained productive motility moved as fast as controls, suggesting that tralopyril primarily reduces the proportion of parasites capable of sustaining productive motility. Consistent with the effects seen here, Portwood *et al*. reported that tralopyril dramatically impaired *P. berghei* and *P. falciparum* development, affecting ookinete and sporozoite motility, oocyst formation, and sporozoite maturation, likely by uncoupling mitochondrial respiration and impairing ATP production (Portwood, 2026). These observations support the emerging concept that certain vector control compounds can exert direct effects on the parasite in addition to their insecticidal activity.

Hydramethylnon and cyflumetofen, a mitochondrial complex III (Song & Scharf, 2009) and II inhibitor (Hayashi, 2012) respectively, produced a modest reduction in motility. Taken together with the impacts of tralopyril, this suggest that even slight perturbations in the electron transport chain may impair sporozoite gliding motility (van Dooren, Stimmler, & McFadden, 2006).

Clodinafop propargyl, an acetyl-CoA inhibitor involved in fatty acid biosynthesis (Secor & Cseke, 1988), also showed a trend towards reduced sporozoite motility. Sporozoites rely on continuous membrane turnover during gliding motility (Frischknecht & Matuschewski, 2017; Munter et al., 2009); hence, interference with lipid biosynthesis could subtly impair membrane dynamics or membrane secretion, processes that are essential for organelle biogenesis and vesicular trafficking in apicomplexan parasites (Coppens, Sinai, & Joiner, 2000; Frischknecht & Matuschewski, 2017; van Dooren et al., 2006). Indeed, clodinafop-propargyl has been reported to inhibit growth in *Toxoplasma gondii* (Zuther, Johnson, Haselkorn, McLeod, & Gornicki, 1999). These findings highlight lipid metabolism and mitochondrial respiration as potential vulnerabilities in the mosquito parasite stages.

Across compound classes, the combined analysis of productive motility and migration speed reveals distinct modes of action. Avermectins act as potent binary inhibitors that stop motility, mitochondrial inhibitors reduce the proportion of parasites capable of motility without significantly altering movement dynamics, and pyrethroids impair the kinematics of motility in parasites that remain active. Sporozoite speed is highly regulated and adapted to evade host immune responses (Amino et al., 2006; Ejigiri et al., 2012; Hellmann et al., 2011), and deviations may potentially impair tissue traversal and transmission success (Hellmann et al., 2011). This distinction highlights that successful transmission can be disrupted through multiple mechanistic routes, including complete inhibition of motility, reduced likelihood of motility initiation, or impaired movement efficiency.

In conclusion, our study reveals that dual vector-parasite activity amongst insecticidal compounds may be far more common than previously appreciated. 15% of the compounds screened impaired *Plasmodium* sporozoite motility, highlighting the parasite as an unanticipated target of several existing insecticides. Whilst mitochondrial inhibitors emerged as a particularly strong class affecting parasite transmission stages, we also identified pronounced effects for avermectins, neurotoxic insecticides whose primary targets lie within the mosquito nervous system. These findings demonstrate that compounds developed to kill mosquitoes can simultaneously disrupt parasite biology through distinct mechanisms and should be tested for such activities. Thus, incorporating transmission blocking assays into vector control discovery pipelines may therefore reveal previously overlooked transmission-blocking properties of candidate compounds. Such dual-activity insecticides could provide a powerful strategy to counteract resistance, as partial loss of mosquito lethality may be offset by reduced parasite transmission. Systematic evaluation of parasite susceptibility should therefore be considered as a component of next-generation vector control development.

## Materials and methods

### Mosquito rearing

Presumed mated adult female mosquitoes of the species *Anopheles stephensi*, fully susceptible to insecticides (SDA 500, MRA-1326, originally contributed by Peter F. Billingsley from Sindh Province, Pakistan), were used in all experiments. Mosquitoes were reared at Heidelberg University under standard insectary conditions, with an ambient temperature of 27-28°C, 70-80% humidity and a 12:12 h photoperiod with a 1 h dawn:dusk cycle. Larvae were reared in milli-Q water, supplemented with 0.1% salt, and fed twice a day with Tetramin® fish food, whilst adults were fed 10% sucrose *ad libitum* from emergence. Colony maintenance was ensured through regular blood feeding using a Hemotek feeder (Hemotek Ltd, UK) supplied with reconstituted human blood.

### Insecticide dose determination

A stock solution of the insecticide abamectin was prepared in acetone at 10 mg/ml, equivalent to 1%, and diluted to target concentrations. Mosquitoes were topically exposed 3-5 days post-emergence to eight concentrations ranging between 0.0005 % and 1 %, along with an acetone-only control, by applying 0.5 μl of compound dissolved in acetone directly onto the dorsal thorax cuticle with a Hamilton® gastight syringe, as previously described (Brito-Sierra, Kaur, & Hill, 2019; Burgess, King, & Geden, 2020). Mosquitoes were anaesthetised on ice during exposures and subsequently transferred into cups and maintained under standard insectary conditions. Mortality was scored 24 h post-exposure and every subsequent 24 h up to 72 h. For each concentration, 20 mosquitoes were exposed and the experiment was performed in two to four biological replicates. Data were fitted to a sigmoidal four-parameter logistic (4PL) model in GraphPad Prism 10 (GraphPad Software, La Jolla, CA, USA) and lethal concentrations calculated.

### Mouse infections with *P. berghei*

Female CD1 Swiss mice were infected with cryo-preserved stabilates of csGFP *P. berghei*-infected blood (Natarajan et al., 2001) via intraperitoneal injection and, upon parasitaemia of ~2%, a full blood transfer with 20 million parasites was performed under anaesthesia (120 mg/kg ketamine, 16 mg/ml xylazine) into a second, naïve mouse. Four days post-transfer, exflagellation was assessed *in vitro* at 21°C.

### *An. stephensi* infections with *P. berghei*

Upon sufficient exflagellation, mice were anaesthetised and offered to 3-5-day old female *An. stephensi* mosquitoes as a blood meal. Infectious feeds were performed at 21°C and 80% humidity, non-bloodfed mosquitoes were removed approximately 24 h post-infection, and the bloodfed mosquitoes maintained under the same conditions throughout the experiment.

### Topical exposure of *P. berghei*-infected *An. stephensi* with abamectin and hydramethylnon

*P. berghei*-infected adult female mosquitoes were topically exposed to abamectin and hydramethylnon 10 days post-infection by applying 0.5 μl of compound dissolved in acetone directly onto the dorsal thorax cuticle with a gastight syringe (Hamilton Company, Switzerland), whilst an untreated control group was exposed to acetone only, as previously described (Lees et al., 2019). Mosquitoes were anaesthetised on ice during exposures. Exposed mosquitoes were subsequently maintained at 21°C and 80% humidity upon infection for 72 h prior to dissections. For hydramethylnon, a separate exposure was performed 30 min post-infection as described above until dissections five days post-infection and exposure.

### Oocyst counts and measurements

At three days post-exposure and 13-days post-infection, or five days post-exposure and infection for early hydramethylnon-exposed mosquitoes, midguts were dissected in phosphate-buffered saline (PBS; Sigma Aldrich, Germany), permeabilised for 20 min in 2% Ninodet-P40 (NP40; Biozol, Germany)/PBS, and stained with 0.1% mercurochrome (Sigma Aldrich, Germany)/PBS for 30 min. Midguts were then imaged at 10x magnification on a Zeiss Axio Lab.A1 microscope. Prevalence was determined based on presence of oocysts. Oocysts were quantified and their area measured using ImageJ (version 1.54p) (Schneider, Rasband, & Eliceiri, 2012). Intact and burst oocysts were distinguished based on morphology and contrast; both were included in total oocyst counts, while only intact oocysts were used for area measurements. In each case a minimum of three independent infections were used. Significant differences between control and insecticide-exposed groups were determined using a Fisher’s exact test for prevalence and a Mann-Whitney test or one-way ANOVA with multiple comparisons for oocyst counts and percentage burst oocysts in GraphPad Prism 10 (GraphPad Software, La Jolla, CA, USA), with a p-value ≤0.05 considered statistically significant.

### Sporozoite counts

At the same timepoint as for oocyst assessment, sporozoites were collected from the haemolymph and salivary glands. Haemolymph was obtained via perfusion: the last abdominal segment was removed and small volumes (5-10 μl) of PBS were injected into the thorax, with the resulting flow-through collected. Samples were pooled from up to 20 mosquitoes. Prevalence of salivary gland sporozoites in csGFP *P. berghei* infections was determined based on GFP signal, expressed by the contained sporozoites under the CSP promoter (Natarajan et al., 2001). Salivary glands were dissected into 20 μl PBS in pools of up to five mosquitoes and manually homogenised for two min to release the contained sporozoites. For both haemolymph and salivary gland samples, sporozoites were counted using a haemocytometer at 40x magnification on a Zeiss Axio Lab.A1 microscope (Zeiss, Germany). Total sporozoite numbers per sample were extrapolated and normalised to the number of mosquitoes to obtain the average sporozoite count per mosquito. Significant differences between control and insecticide-exposed groups were assessed using a Fisher’s exact test for prevalence and a Mann-Whitney test or one-way ANOVA with multiple comparisons for sporozoite counts in GraphPad Prism 10 (GraphPad Software, La Jolla, CA, USA), with a p-value ≤0.05 considered statistically significant.

### *In vitro* sporozoite motility assays

100-fold stock solutions of each tested compound were prepared in dimethyl sulfoxide (DMSO; Sigma Aldrich, Germany) and diluted to two-fold stock solutions in RPMI 1640 medium (RPMI; Corning, Manassas, VA, USA) for final concentrations of 1% DMSO and 9 µg/ml, 13.5 µg/ml and 22.5 µg/ml of compound, reflecting the concentrations expected in mosquitoes after contact with pyrethroid-treated surfaces (Spielmeyer et al., 2019). A DMSO-only control was additionally prepared in parallel. *In vitro* motility assays were conducted as previously described (Prinz et al., 2017). In brief, infected, unexposed *An. stephensi* were used 17-21 days post-infection. Salivary glands from 5-20 mosquitoes were dissected into cold RPMI, the sporozoites were mechanically extracted and diluted in cold RPMI as needed to yield a total of 100-200 sporozoites per field of view during the assay. Sporozoites were activated with 12% bovine serum albumin (BSA; Sigma Aldrich, Germany)/RPMI and combined in a 1:1 ratio with the two-fold insecticide stocks in a 96 well optical plate (Thermo Scientific, Germany), yielding the final desired compound concentrations and 3% BSA. Following centrifugation to settle sporozoites at the bottom of each well, they were imaged at 25x magnification in DIC and GFP on an inverted Zeiss Axiovert 200M microscope (Zeiss, Germany) at 0 min, 10 min and 60 min after compound addition. For each concentration and time point, sporozoites were recorded for 3 min at 20 frames per minute in either filter. Z projections of each video were generated in ImageJ (version 1.54p) (Schneider et al., 2012), and motility patterns were classified as productive (circular) or non-productive (non-circular). Three biological replicates were performed for each compound.

For *in vivo* abamectin exposure, additional concentrations of 0.9 µg/ml, 2.7 µg/ml, 3.6 µg/ml, 4.5 µg/ml, 5.4 µg/ml and 7.2 µg/ml were tested to provide a finer dose-response analysis. Significant differences in productive motility compared to the control were determined using a two-way ANOVA with multiple comparisons in GraphPad Prism, with a p-value ≤0.05 considered significant.

### *Ilastik*-based automated sporozoite motility tracking and labelling

To automate the analysis of large datasets of recordings from *in vitro* sporozoite motility assays and reduce human bias, a machine-learning workflow was established. The backbone was based on *Ilastik* (v1.3.3) (Sommer C., 2011), trained on manual annotations from a representative subset. For each recording, the workflow yielded two complementary outputs: (i) object-level phenotype quantification (e.g., the fraction of sporozoites exhibiting rotational versus non-rotational behaviour per compound and condition) and (ii) an estimate of rotational motility computed for the subset of rotating sporozoites per compound and condition.

#### Data pre-processing

To reduce computational burden and manual annotation effort, all microscopy recordings were pre-processed using spatial down sampling followed by temporal projection.

#### Spatial down sampling

Each frame was down sampled by a factor of four in both spatial dimensions using 4×4 max pooling. This reduced the total number of pixels per frame by a factor of 16. Max pooling was selected over average pooling because it preferentially preserves bright, thin structures characteristic of sporozoites, thereby maintaining relevant morphological features while reducing data size.

#### Temporal projection

Each recording consisted of a time series of *K* frames *I*_*k*_ (*y,x*) where *k* ∈ {1, …, *K*} indexed time and (*x,y*) denoted pixel coordinates. A stable two-dimensional projection image was computed as the temporal mean of the spatially down sampled frames,

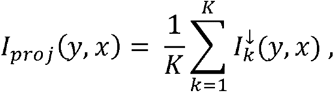

where 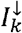 denotes the downsampled frame at time *k*.

The projection yielded a single 2D image that enhanced the visual distinction between rotational and non-rotational phenotypes: rotating sporozoites appeared as compact, approximately circular traces, whereas non-rotating (stationary or translational) sporozoites retained an elongated morphology without circular accumulation (see Fig. 1 for a visual example).

#### Output data structure

For each recording, pre-processing produced:

I. A projection-only 2D image *I*_*proj*_ for robust object detection and phenotype quantification (fraction rotating vs. non-rotating per condition),
II. A multi-slice stack with slice 0 corresponding to *I*_*proj*_ and slices 1..*K* contained the down sampled frames 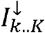

This structure preserved the full temporal information required for estimating rotational motility while providing a stable reference image for segmentation.

#### Ilastik-based segmentation and object classification

For both outcomes (fraction of rotational sporozoites and estimation of rotational motility), an *Ilastik*-based supervised machine-learning pipeline was employed. The pipeline consisted of two sequential *Ilastik* stages with distinct roles: first, pixel-wise segmentation was used to separate sporozoite pixels from background; second, object classification detected individual sporozoites in both the 2D projection and the time-stack representation.

In total, four *Ilastik*-based classifiers were trained on manually annotated data and subsequently applied for automated processing. *Ilastik* is based on a Random Forest classifier (Breiman, 2001) trained from sparse user annotations (Berg et al., 2019; Sommer C., 2011). For pixel classification, each pixel is represented by a set of multi-scale image features derived from the raw input, including intensity- and colour-based measurements, edge/gradient information, and texture descriptors obtained from image filters. Training was performed interactively by placing brush-stroke annotations for both classes on representative recordings and iteratively refining annotations until predictions were stable across multiple recordings.

For object classification, connected parasite instances were extracted from the pixel-class probability maps using *Ilastik’s* connected-component analysis. Each object was represented by an object-level feature vector (intensity statistics and geometric/shape descriptors), and a supervised Random Forest classifier was trained by manually labelling a subset of objects. During inference, the classifier produced phenotype predictions for all detected objects. For the stacked representation (slice 0 projection + slices 1..*K* frames), object identity maps were additionally exported over time, enabling tracking of individual sporozoites and quantification of rotational events.

#### Automated processing

Automated analysis of sporozoite motility was performed using trained *Ilastik* models to allow consistent processing of large datasets without manual intervention. For each recording, pixel-wise classification produced probability maps *P*_*c*_ (*y,x*) distinguishing sporozoite pixels from background, which were subsequently used together with the raw images for object classification. This stage generated object prediction tables with phenotype labels for each detected object, and, for rotational motility analysis, per-frame object identity maps were exported to track individual sporozoites over time.

For quantification of non-rotating sporozoites, the fraction of objects assigned to each phenotype class *c* was computed as

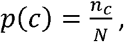

where *N* is the total number of detected objects and *n*_*c*_ the number of objects labelled as class *c*. Two phenotypes were considered: rotating and non-rotating sporozoites, based on the temporally averaged 2D representation, and the primary outcome measure was the fraction of non-rotating objects per recording.

Rotational motility was assessed for reliably segmented objects exhibiting clear rotational behaviour. For each frame *t* of the multi-stack recording (excluding the projection slice at *t* =0) the centre of the object mask was calculated as

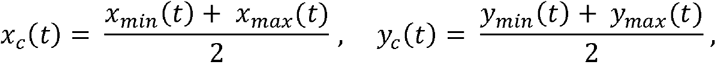

with objects missing in any frame excluded from analysis. An angular signal was constructed for each object as

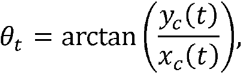

and slow baseline drift was removed using a running mean over a fixed window of *w*= 4 frames:

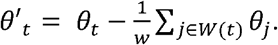

To reduce measurement noise, the drift-corrected angle signal was further smoothed using a Kalman filter, which models angular velocity as approximately constant to capture true rotational trends.

Rotation was quantified by counting sign changes in the drift-corrected signal:

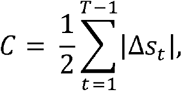

where

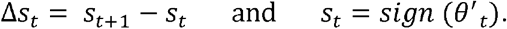

Rotation was computed both from the drift-corrected and Kalman-smoothed signals, and estimates were accepted if the relative discrepancy between the two satisfied

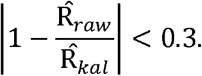

For accepted objects, the final rotation estimate was the average of the two measures,

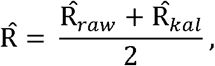

and values were averaged across selected objects within a recording, then across replicates for each experimental condition.

#### Rotation measurement of productively moving sporozoites

To assess the effect of insecticide exposure on the rotation speed of productively motile sporozoites, we performed linear mixed-effects modelling on individual sporozoite rotation counts. The analysis was restricted to sporozoites exhibiting productive motility, as determined by the automated image analysis pipeline. For each compound-concentration-timepoint combination, individual rotation values from all three experimental replicates were pooled and analysed using a linear mixed-effects model (LMM) with the *lme4* (Bates, 2015) and *lmerTest* (Kuznetsova, 2017) packages in R (version 2024.09.0, Build 375).

The model structure was: rotations ~ concentration × timepoint + (1|replicate), where concentration was treated as a categorical factor with the control condition (0 µg/ml) as the reference level, and replicate was included as a random intercept to account for the nested structure of sporozoites within experimental replicates. For each compound, pairwise contrasts between each concentration and the control were extracted using the *emmeans* package (Lenth, 2026), yielding nine comparisons per compound (three concentrations × three timepoints). Raw p-values were corrected for multiple testing using the Benjamini-Hochberg false discovery rate (FDR) procedure across all tests. Statistical significance was defined as FDR < 0.05.

### P. falciparum gametocyte culturing

NF54 *P. falciparum* asexual parasites (provided by Teun Bousema, Radboud University Medical Center, Nijmegen, Netherlands) were cultured in human O+ erythrocytes (Blood bank, University Hospital Heidelberg, Germany) at 2.5% haematocrit and maintained in a medium consisting of RPMI supplemented with 25 mM HEPES, 10% heat-inactivated human serum (Haema AG, Leipzig, Germany), 10 mg/L hypoxanthine (Sigma Aldrich, Germany) and 0.3% sodium bicarbonate (ThermoFisher Scientific, Germany). Cultures were maintained at 37°C under a gas mixture of 5% CO_2_, 5% O_2_ and 90% N_2_. Gametocyte development was promoted by diluting parasite cultures to 0.2% parasitaemia at 5% haematocrit, with a daily medium change over 16-18 consecutive days, whilst avoiding disruption of the erythrocyte layer.

### An. stephensi infections with P. falciparum

0.4% stage V gametocytaemia was achieved through counting thin blood smears and dilution with reconstituted human O+ blood and fed to 3–5-day old mosquitoes in glass membrane feeders pre-warmed to 37°C. Mosquitoes were then maintained in humidified incubators under standard insectary conditions at 28°C. At 24 h post-infection, non-bloodfed and dead mosquitoes were anaesthetised on ice and removed.

### Topical exposures with abamectin of *P. falciparum*-infected *An. stephensi*

*P. falciparum*-infected adult female mosquitoes were topically exposed to abamectin by applying 0.5 μl of compound dissolved in acetone directly onto the dorsal thorax cuticle with a Hamilton® gastight syringe, while an untreated control group was exposed to acetone only, as previously described (Lees et al., 2019). Mosquitoes were anaesthetised on ice during exposures. Exposed mosquitoes were maintained at 21°C under otherwise standard insectary conditions upon infection for 72 h prior to killing and dissections.

### Parasite development analysis

Oocyst counts and measurements, as well as haemolymph and salivary gland sporozoite counts, were performed as described for *P. berghei*. Sporozoites from salivary glands and haemolymph were pooled in groups of 2-20 individuals and extrapolated to an average sporozoite number per mosquito.

## Acknowledgements

This project was funded by the Deutsche Forschungsgemeinschaft (DFG, German Research Foundation), project no. 240245660 – SFB 1129, Gates Foundation (INV-050587), Deutsches Zentrum für Infektionsforschung (DZIF, TTU03.705) and ERC Starting Grant (Project Number 101075635, ReMVeC) to VAI. We would like to thank Dr. David Hong for his scientific counsel throughout the study, Dr. Harsha Vardan Reddy for assisting in the sporozoite motility assay optimisation and Dr. Julia Sattler for assisting with administrative tasks regarding mouse work. We are grateful to Jakob Kerbl-Knapp and Sina Kühnel for their help in rearing testing mosquitoes and Patrick Hörner for assisting with mouse work. Graphs were created in GraphPad Prism and figures with BioRender.

## Supplementary materials

### Supplementary Tables

**Supplementary Table 1. Insecticide overview for sporozoite motility assays**.

**Supplementary Table 2. Sporozoite motility ratios**.

**Supplementary Table 3. Abamectin sporozoite motility dose response**.

**Supplementary Table 4. Sporozoite speed dynamics**.

### Supplementary Figures

**Supplementary Figure 1.**
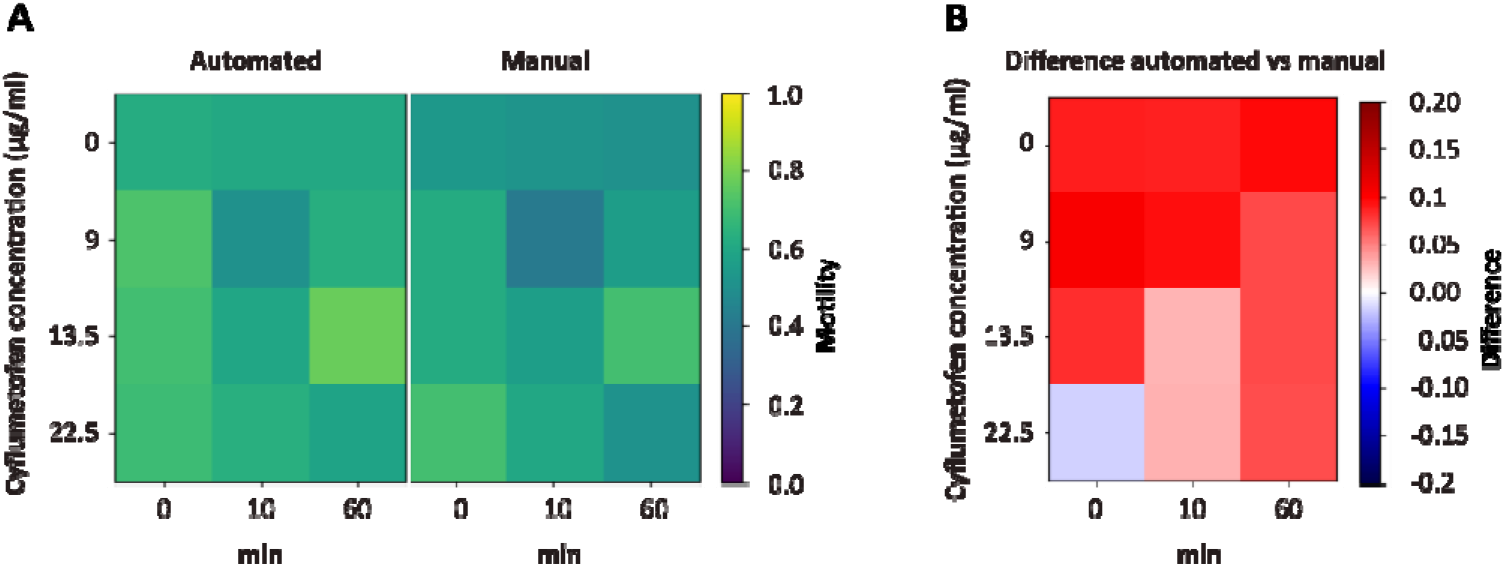
*Ilastik*-based automatic tracking and labelling of *P. berghei in vitro* sporozoite motility data. **(A)** Comparison of automated vs. manual labelling accuracy. The error between automated and manual labelling is represented for cyflumetofen, with the heatmaps showing the productive motility for 0, 9, 13.5, and 22.5 µg/ml at timepoints 0-, 10, and 60-min post addition of the compound on a scale of 0 - 1 (blue to yellow), with yellow = increased and blue = reduced productive motility (left). **(B)** The difference between the automated vs. manual labelling is presented as a heatmap, with >0 = larger difference (red) and <0 = smaller difference (blue).

**Supplementary Figure 2.**
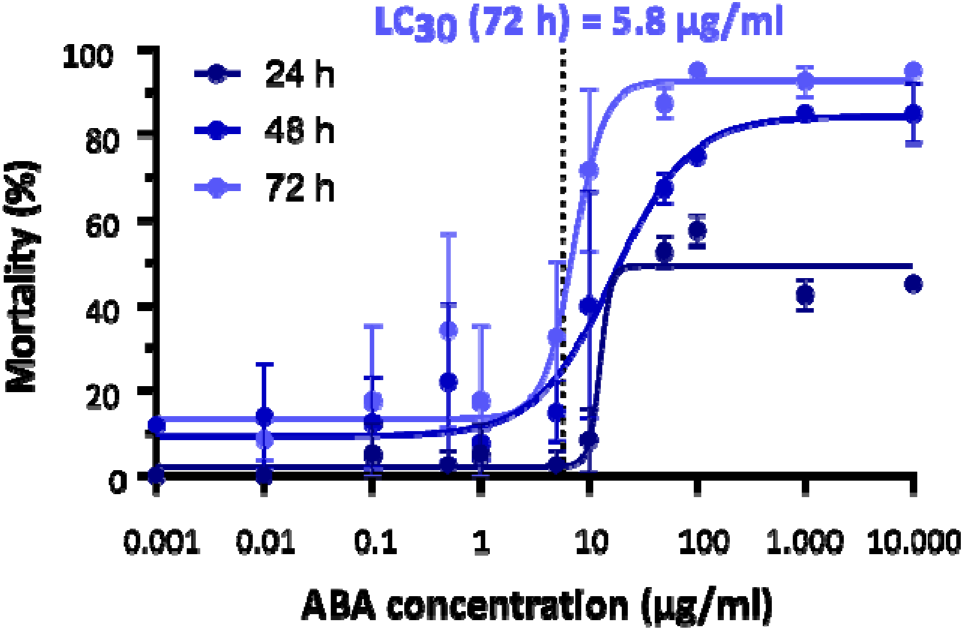
Abamectin dose response following topical exposure. Percent mortality per concentration (µg/ml) abamectin (ABA) dissolved in acetone at 24 h (dark blue), 48 h (medium blue), and 72 h (light blue) post-topical exposure at 21 °C. LC_30_ at 72 h indicated by dashed line. Data points represent means and error bars standard deviation. 20 mosquitoes per replicate, 2-4 replicates per concentration.

**Supplementary Figure 3.**
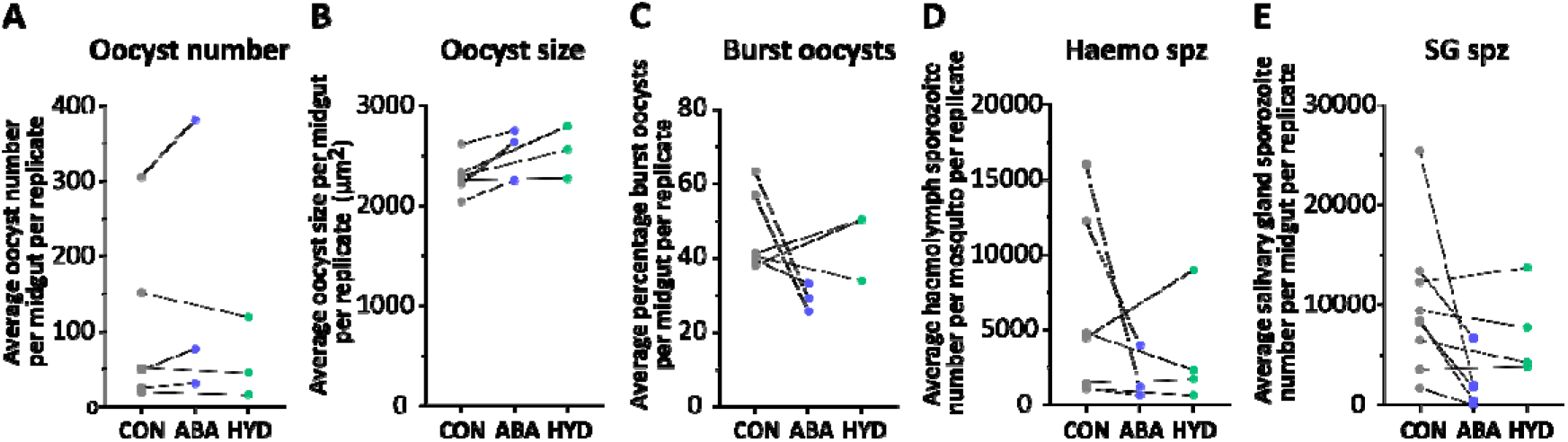
Effects of *in vivo* abamectin and hydramethylnon exposure on *P. berghei*-infected mosquitoes per replicate. **(A-E)** *P. berghei* oocyst and sporozoite counts, as well as burst oocyst percentage, per replicate for control (CON, grey) vs. abamectin-(ABA, blue) and hydramethylnon-exposed (HYD, green) mosquitoes 13 days post-infection and three days post-exposure. **(A)** Mean oocyst number per midgut. **(B)** Mean average oocyst size per midgut in µm^2^ per midgut. **(C)** Mean percentage burst oocysts per midgut. **(D)** Mean haemolymph sporozoite number per mosquito. **(E)** Mean salivary gland sporozoite number per mosquito.

**Supplementary Figure 4.**
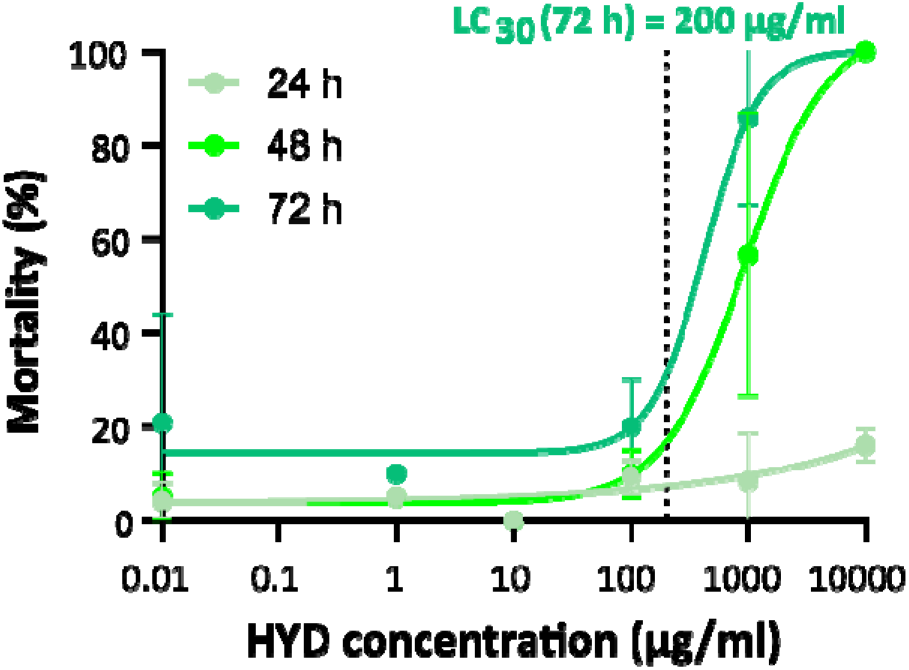
Hydramethylnon dose response following topical exposure. Percent mortality per concentration (µg/ml) hydramethylnon (HYD) dissolved in acetone at 24 h (light green), 48 h (medium green), and 72 h (dark green) post-topical exposure at 21 °C. LC_30_ at 72 h indicated by dashed line. Lines represent means and error bars standard deviation. 20 mosquitoes per replicate, 2-4 replicates per concentration.

**Supplementary Figure 5.**
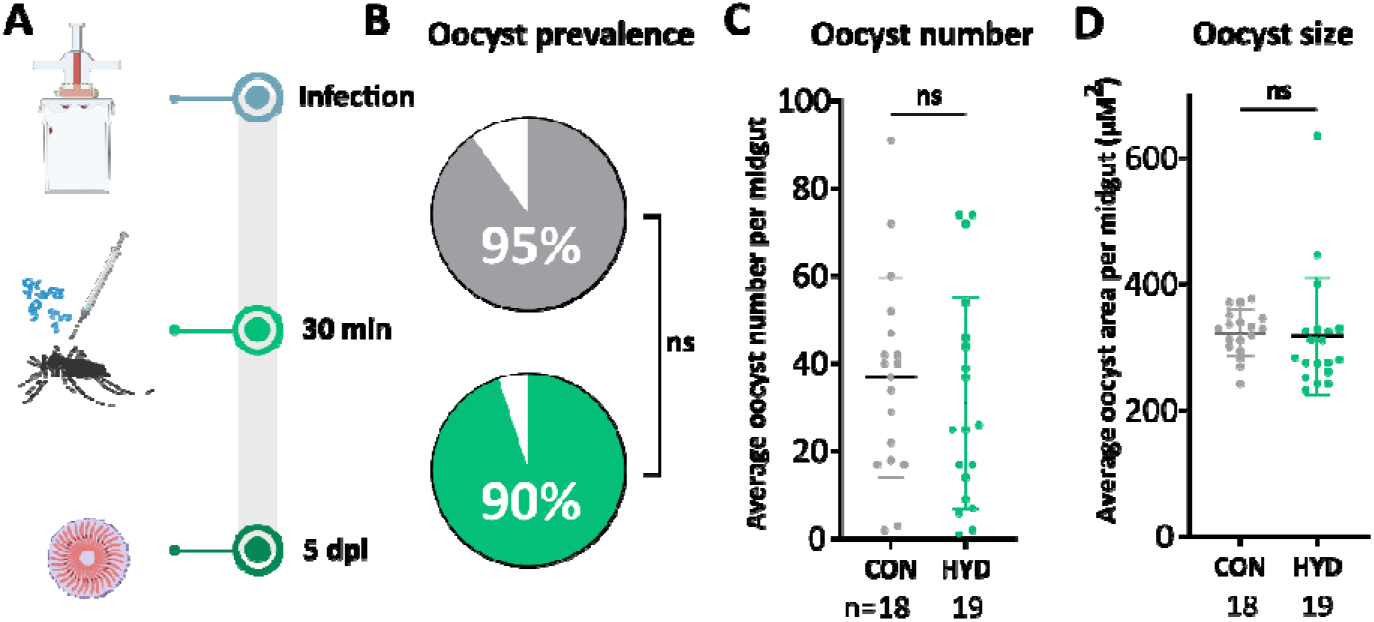
*P. berghei* infection responses to *in vivo* hydramethylnon exposure. **(A)** Schematic of *in vivo* hydramethylnon exposure of *P. berghei*-infected mosquitoes for late early oocyst (right) development assessment. Mosquitoes were topically exposed to the LC_30_ concentration of hydramethylnon at 72 h (200 µg/ml) or an acetone control 30 min post-infection and dissected five days post-infection and exposure for oocyst assessment. **(B-D)** *P. berghei* infection responses to hydramethylnon (HYD, green) vs control (CON, grey) exposure five days post-infection and exposure. Significant differences in prevalence determined by Fisher’s exact test and for oocyst measurements by Mann-Whitney test, with p<0.05 considered significant. Lines represent means and error bars indicate standard deviation. **(B)** Oocyst prevalence five days post-infection and exposure (n CON= 20, HYD = 20). **(C)** Mean oocyst number five days post-infection and exposure (n CON = 18, HYD = 19). **(D)** Mean oocyst size (µm^2^) five days post-infection and exposure (n CON = 18, HYD = 20).

**Supplementary Figure 6.**
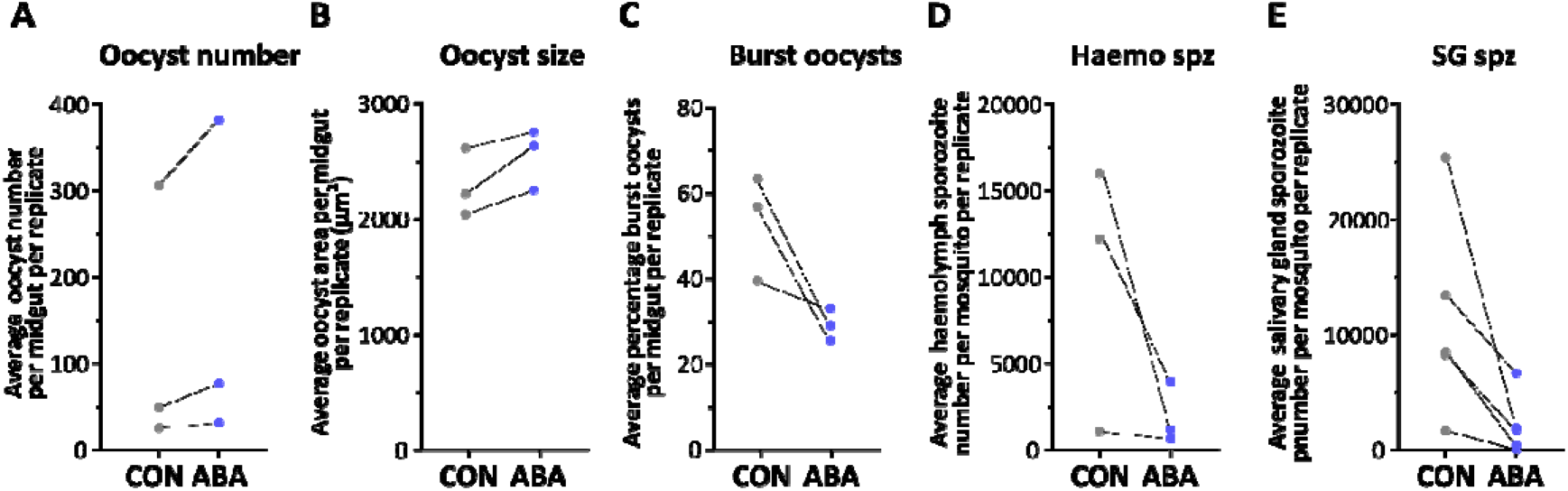
Effects of *in vivo* abamectin exposure of *P. falciparum*-infected mosquitoes per replicate. **(A-E)** *P. falciparum* oocyst and sporozoite counts, as well as burst oocyst percentage, per replicate for control (CON, grey) vs. abamectin-exposed (ABA, blue) mosquitoes 13 days post-infection and three days post-exposure. **(A)** Mean oocyst number per midgut. **(B)** Mean average oocyst size per midgut in µm per midgut. **(C)** Mean percentage burst oocysts per midgut. **(D)** Mean haemolymph sporozoite number per mosquito. **(E)** Mean salivary gland sporozoite number per mosquito.

