## Supplementary Table 1 for "Insecticides can simultaneously target mosquito vectors and malaria parasites"

Supplementary Table 1: Compound list with IRAC information

| Compound | Insecticide Resistance Action Committee (IRAC) Group | IRAC Subgroup | Notes |
|---|---|---|---|
| Abamectin | Glutamate-gated chloride channel (GluCl) allosteric modulators | Avermectins, milbemycins | |
| Acifluorfen | N/A | N/A | |
| Alpha-cypermethrin | Sodium channel modulator | Pyrethroids, pyrethrins | |
| Bendiocarb | Acetylcholinesterase (AchE) inhibitors | Carbamates | |
| Bifenthrin | Sodium channel modulator | Pyrethroids, pyrethrins | |
| Broflanilide | GABA-gated chloride channel allosteric modulators | Meta-diamides, isoxazolines | |
| Chlorfenapyr | Uncouplers of oxidative phosphorylation via disruption of the proton gradient | Pyrroles, dinitrophenols, sulfluramid | |
| Clodinafop-propargyl | N/A | N/A | Acetyl-CoA-carboxylase inhibitor, herbicide |
| Clothianidin | Nicotinic acetylcholine receptor (NAchR) competitive modulators | Neonicotinoids | |
| Cyflumetofen | Mitochondrial complex II electron transport inhibitors | Beta-ketonitrile derivatives | |
| Deltamethrin | Sodium channel modulator | Pyrethroids, pyrethrins | |
| Diafenthiuron | Inhibitors of mitochondrial ATP synthase | Diafenthiuron | |
| Diflubenzuron | Inihibitors of chitin biosynthesis affecting CHS1 | Benzoylureas | |
| DNOC | Uncouplers of oxidative phosphorylation via disruption of the proton gradient | Pyrroles, dinitrophenols, sulfluramid | |
| Emamectin-benzoate | Glutamate-gated chloride channel (GluCl) allosteric modulators | Avermectins, milbemycins | |
| Etofenprox | Sodium channel modulator | Pyrethroids, pyrethrins | |
| Fenpyroximate | Mitochondrial complex I electron transport inhibitors | Meti acaricides and insecticides | |
| Flufenoxuron | Inihibitors of chitin biosynthesis affecting CHS1 | Benzoylureas | |
| Hydramethylnon | Mitochondrial complex III electron transport inhibitors – QO site | Hydramethylnon | |
| Imidacloprid | Nicotinic acetylcholine receptor (NAchR) competitive modulators | Neonicotinoids | |
| Ivermectin | Glutamate-gated chloride channel (GluCl) allosteric modulators | Avermectins, milbemycins | |
| Malaoxon | N/A | N/A | Active metabolite of malathion |
| Novaluron | Inihibitors of chitin biosynthesis affecting CHS1 | Benzoylureas | |
| Permethrin | Sodium channel modulator | Pyrethroids, pyrethrins | |
| Piperonyl butoxide | N/A | N/A | Synergist |
| Pirimiphos-methyl | Acetylcholinesterase (AchE) inhibitors | Organophosphates | |
| Pyriproxyfen | Juvenile hormone receptor modulators | Pyriproxyfen | |
| Shelock | N/A | N/A | Mitochondrial complex I inhibitor, unlicensed |
| Spinosad | Nicotinic acetylcholine receptor (NAchR) allosteric modulators – Site I | Spinosyns | |
| Teflubenzuron | Inihibitors of chitin biosynthesis affecting CHS1 | Benzoylureas | |
| Tralopyril | N/A | N/A | Active metabolite of chlorfenapyr |
| Transfluthrin | Sodium channel modulator | Pyrethroids, pyrethrins | |
