## Supplementary Table 2 for "Insecticides can simultaneously target mosquito vectors and malaria parasites"

Supplementary Table 2: Raw data and statistics from motility assessment

| Compound | Mode of action | Time | 9 µg/ml ratio | 9 µg/ml p-value | 13.5 µg/ml ratio | 13.5 µg/ml p-value | 22.5 µg/ml ratio | 22.5 µg/ml pval |
|---|---|---|---|---|---|---|---|---|
| Abamectin | Avermectin | 0 min | 0.036 | 0.0001 | 0.012 | 0.0124 | 0.003 | 0.0001 |
| | | 10 min | 0.014 | 0.0001 | 0.004 | 0.0001 | 0.001 | 0.0001 |
| | | 60 min | 0.007 | 0.0001 | 0.002 | 0.0001 | 0 | 0.0001 |
| Acifluorfen | itochondrial inhibit | 0 min | 0.918 | 0.9465 | 0.933 | 0.9689 | 0.94 | 0.9774 |
| | | 10 min | 1.04 | 0.9983 | 1.111 | 0.9677 | 0.812 | 0.867 |
| | | 60 min | 1.163 | 0.8689 | 1.017 | 0.9998 | 0.843 | 0.8801 |
| Alpha-cyp | Pyrethroid | 0 min | 1.445 | 0.6908 | 1.273 | 0.8694 | 1.411 | 0.5029 |
| | | 10 min | 0.728 | 0.9898 | 0.73 | 0.9386 | 0.588 | 0.9999 |
| | | 60 min | 0.749 | 0.9925 | 0.83 | 0.9845 | 1.183 | 0.6265 |
| Bendiocarb | Carbamate | 0 min | 0.828 | 0.9905 | 1.182 | 0.6953 | 0.737 | 0.8013 |
| | | 10 min | 0.896 | 0.9975 | 0.931 | 0.992 | 0.929 | 0.9677 |
| | | 60 min | 0.64 | 0.93 | 1.586 | 0.9073 | 2.653 | 0.4785 |
| Bifenthrin | Pyrethroid | 0 min | 1.429 | 0.2123 | 1.551 | 0.1241 | 1.738 | 0.0559 |
| | | 10 min | 0.947 | 0.779 | 0.629 | 0.6628 | 0.672 | 0.7462 |
| | | 60 min | 0.82 | 0.9697 | 1.235 | 0.5892 | 0.921 | 0.9006 |
| Broflanalide | GABA modulator | 0 min | 1.063 | 0.9883 | 1.21 | 0.9977 | 0.603 | 0.3273 |
| | | 10 min | 1.309 | 0.9633 | 0.851 | 0.9931 | 1.02 | 0.9977 |
| | | 60 min | 0.961 | 0.9999 | 0.626 | 0.6666 | 0.659 | 0.8398 |
| Chlorfenapyr | New-net | 0 min | 0.606 | 0.9456 | 0.945 | 0.9808 | 1.131 | 0.8741 |
| | | 10 min | 0.77 | 0.5155 | 1.572 | 0.2697 | 1.379 | 0.7573 |
| | | 60 min | 0.482 | 0.1805 | 0.607 | 0.5076 | 0.594 | 0.5012 |
| Clodinafop-propargy | ACC inhibitor | 0 min | 1.103 | 0.8676 | 1.3 | 0.3539 | 1.182 | 0.7727 |
| | | 10 min | 0.665 | 0.5894 | 0.733 | 0.8178 | 0.68 | 0.5958 |
| | | 60 min | 0.338 | 0.257 | 0.541 | 0.3388 | 0.651 | 0.4265 |
| Clothianidin | Neonicotinoid | 0 min | 0.886 | 0.9829 | 1.136 | 0.9959 | 1.23 | 0.9949 |
| | | 10 min | 0.871 | 0.9807 | 0.989 | 0.9975 | 1.028 | 0.9564 |
| | | 60 min | 1.225 | 0.9872 | 2.2 | 0.8295 | 0.989 | 0.9994 |
| Cyflumetofen | itochondrial inhibit | 0 min | 0.677 | 0.6646 | 0.644 | 0.6492 | 0.476 | 0.4711 |
| | | 10 min | 0.94 | 0.9997 | 0.564 | 0.5437 | 0.425 | 0.3874 |
| | | 60 min | 0.739 | 0.4834 | 0.51 | 0.0571 | 0.837 | 0.7222 |
| Deltamethrin | Pyrethroid | 0 min | 1.752 | 0.1375 | 2.052 | 0.0607 | 1.961 | 0.0598 |
| | | 10 min | 1.019 | 0.9549 | 0.768 | 0.9999 | 0.759 | 0.8373 |
| | | 60 min | 0.909 | 0.9887 | 0.937 | 0.9957 | 0.874 | 0.9991 |
| Diafenthiuron | itochondrial inhibit | 0 min | 1.013 | 0.999 | 0.927 | 0.9998 | 0.788 | 0.8232 |
| | | 10 min | 0.762 | 0.7572 | 0.941 | 0.9029 | 0.658 | 0.4427 |
| | | 60 min | 0.775 | 0.982 | 0.944 | 0.9996 | 0.547 | 0.7372 |
| Diflubenzuron | Benzoylurea | 0 min | 0.962 | 0.9606 | 0.967 | 0.9885 | 1.029 | 0.9863 |
| | | 10 min | 0.707 | 0.6053 | 0.882 | 0.4315 | 0.599 | 0.3414 |
| | | 60 min | 0.896 | 0.9463 | 1.005 | 0.9528 | 1.073 | 0.9848 |
| DNOC | itochondrial inhibit | 0 min | 0.894 | 0.9819 | 0.91 | 0.9966 | 1.275 | 0.6625 |
| | | 10 min | 1.516 | 0.5031 | 1.063 | 0.9596 | 0.896 | 0.9923 |
| | | 60 min | 0.932 | 0.9999 | 0.735 | 0.7049 | 0.668 | 0.5017 |
| Emamectin-Benzoate | Avermectin | 0 min | 0.854 | 0.9513 | 0.794 | 0.8691 | 0.164 | 0.0018 |
| | | 10 min | 1.157 | 0.7749 | 1.003 | 0.9999 | 0.026 | 0.0001 |
| | | 60 min | 1.124 | 0.9811 | 0.671 | 0.3868 | 0.006 | 0.0001 |
| Etofenprox | Pyrethroid | 0 min | 1.208 | 0.993 | 1.69 | 0.7719 | 1.713 | 0.5113 |
| | | 10 min | 0.946 | 0.9992 | 0.773 | 0.914 | 0.801 | 0.9999 |
| | | 60 min | 0.695 | 0.9657 | 0.58 | 0.8256 | 0.935 | 0.9989 |
| Fenpyroximate | itochondrial inhibit | 0 min | 0.826 | 0.8895 | 0.553 | 0.3551 | 0.482 | 0.2285 |
| | | 10 min | 0.351 | 0.1227 | 0.323 | 0.0931 | 0.336 | 0.0981 |
| | | 60 min | 0.664 | 0.6704 | 0.601 | 0.5408 | 0.671 | 0.6995 |
| Flufenoxuron | Benzoylurea | 0 min | 1.233 | 0.6907 | 0.844 | 0.717 | 0.78 | 0.5086 |
| | | 10 min | 0.857 | 0.9731 | 0.551 | 0.3702 | 0.681 | 0.7178 |
| | | 60 min | 0.61 | 0.7595 | 0.491 | 0.6434 | 0.114 | 0.2074 |
| Hydramethylnon | itochondrial inhibit | 0 min | 1.526 | 0.241 | 1.08 | 0.9648 | 0.726 | 0.4606 |
| | | 10 min | 0.832 | 0.6587 | 0.297 | 0.0027 | 0.042 | 0.0001 |
| | | 60 min | 0.354 | 0.0078 | 0.054 | 0.0005 | 0 | 0.0003 |
| Imidacloprid | Neonicotinoid | 0 min | 0.981 | 0.9999 | 0.802 | 0.8452 | 0.893 | 0.9147 |
| | | 10 min | 0.931 | 0.9976 | 0.811 | 0.9477 | 0.776 | 0.939 |
| | | 60 min | 1.08 | 0.9981 | 0.872 | 0.9992 | 0.531 | 0.5709 |
| Ivermectin | Avermectin | 0 min | 0.013 | 0.0001 | 0.008 | 0.0001 | 0 | 0.0001 |
| | | 10 min | 0.004 | 0.0001 | 0.006 | 0.0001 | 0.001 | 0.0001 |
| | | 60 min | 0.011 | 0.0001 | 0.003 | 0.0001 | 0 | 0.0001 |
| Malaoxoan | Organophosphate | 0 min | 1.078 | 0.9845 | 1.304 | 0.5698 | 1.122 | 0.9452 |
| | | 10 min | 0.854 | 0.9578 | 0.844 | 0.9496 | 0.888 | 0.9805 |
| | | 60 min | 1.024 | 0.9998 | 1.533 | 0.3171 | 1.821 | 0.0761 |
| Novaluron | Benzoylurea | 0 min | 0.859 | 0.7437 | 0.718 | 0.4467 | 0.78 | 0.5444 |
| | | 10 min | 1.141 | 0.9652 | 1.11 | 0.988 | 0.99 | 0.9999 |
| | | 60 min | 0.906 | 0.9051 | 0.941 | 0.9535 | 0.594 | 0.4463 |
| Permethrin | Pyrethroid | 0 min | 2.16 | 0.0404 | 1.421 | 0.3871 | 2.238 | 0.0311 |
| | | 10 min | 1.862 | 0.0413 | 1.855 | 0.0626 | 1.535 | 0.1662 |
| | | 60 min | 0.573 | 0.2709 | 0.958 | 0.9987 | 0.749 | 0.85 |
| Piperonyl-butoxide | New-net | 0 min | 1.218 | 0.9635 | 1.02 | 0.9689 | 1.171 | 0.8198 |
| | | 10 min | 0.747 | 0.5054 | 0.682 | 0.3063 | 0.815 | 0.4716 |
| | | 60 min | 1.294 | 0.8121 | 1.625 | 0.8206 | 1.027 | 0.9986 |
| Pirimiphos-Methyl | Organophosphate | 0 min | 1.653 | 0.4919 | 1.42 | 0.7374 | 1.649 | 0.4954 |
| | | 10 min | 1.07 | 0.9992 | 0.898 | 0.9968 | 1.186 | 0.9872 |
| | | 60 min | 1.646 | 0.8635 | 1.587 | 0.8885 | 1.365 | 0.9634 |
| Pyriproxyfen | New-net | 0 min | 1.215 | 0.9769 | 1.089 | 0.998 | 1.01 | 0.9999 |
| | | 10 min | 0.799 | 0.973 | 0.685 | 0.8942 | 0.746 | 0.9442 |
| | | 60 min | 1.547 | 0.7906 | 1.217 | 0.9749 | 1.393 | 0.8929 |
| Sherlock | itochondrial inhibit | 0 min | 1.104 | 0.9542 | 1.09 | 0.9656 | 0.755 | 0.4125 |
| | | 10 min | 0.671 | 0.8802 | 1.09 | 0.9999 | 0.791 | 0.7718 |
| | | 60 min | 0.754 | 0.9778 | 0.923 | 0.9975 | 0.671 | 0.961 |
| Spinosad | NaCHR modulator | 0 min | 1.029 | 0.9043 | 0.766 | 0.4135 | 0.55 | 0.2522 |
| | | 10 min | 0.863 | 0.9999 | 1.043 | 0.9962 | 0.542 | 0.8031 |
| | | 60 min | 0.91 | 0.0581 | 0.605 | 0.8705 | 0.701 | 0.1396 |
| Teflubenzuron | Benzoylurea | 0 min | 1.175 | 0.9896 | 1.146 | 0.999 | 0.839 | 0.999 |
| | | 10 min | 0.991 | 0.9894 | 1.183 | 0.8424 | 1.284 | 0.841 |
| | | 60 min | 1.001 | 0.8336 | 0.33 | 0.2546 | 0.611 | 0.6527 |
| Tralopyril | New-net | 0 min | 0.699 | 0.8274 | 0.505 | 0.4444 | 0.568 | 0.5771 |
| | | 10 min | 0.286 | 0.0032 | 0.422 | 0.0281 | 0.245 | 0.0015 |
| | | 60 min | 0.065 | 0.0001 | 0.073 | 0.0001 | 0.056 | 0.0001 |
| Transfluthrin | Pyrethroid | 0 min | 1.412 | 0.919 | 2.167 | 0.599 | 1.805 | 0.7758 |
| | | 10 min | 0.963 | 0.9983 | 1.217 | 0.9994 | 1.313 | 0.894 |
| | | 60 min | 1.097 | 0.993 | 1.706 | 0.3363 | 2.032 | 0.1907 |
