## Supplementary Table 3 for "Insecticides can simultaneously target mosquito vectors and malaria parasites"

Supplementary Table 3: LC50 motility response to abamectin

| Abamectin do | 0 min p-value | 10 min p-valu | 60 min p-value |
|---|---|---|---|
| 0.9 | 0.4833 | 0.1339 | 0.8543 |
| 2.7 | 0.7209 | 0.0117 | 0.7049 |
| 3.6 | 0.1878 | 0.8399 | 0.224 |
| 4.5 | 0.1256 | 0.0219 | 0.3754 |
| 5.4 | 0.0165 | 0.0153 | 0.0053 |
| 7.2 | 0.0618 | 0.0106 | 0.004 |
| 9 | 0.0009 | 0.0001 | 0.0002 |
| 13.5 | 0.0001 | 0.0001 | 0.0001 |
| 22.5 | 0.0000 | 0.0000 | 0.0001 |
