## Supplementary Table 4 for "Insecticides can simultaneously target mosquito vectors and malaria parasites"

Supplementary Table 4: Raw data and statistics for rotations in motility screen

| Compound | Timepoint_mi | Concentratior | Estimate | SE | df | t_ratio | p_value_raw | p_adj_FDR | signif_raw | signif_FDR |
|---|---|---|---|---|---|---|---|---|---|---|
| | | 9 | -0.291 | 0.650 | 523.416 | -0.447 | 0.655 | 0.790 | ns | ns |
| | | 13.5 | -0.548 | 0.952 | 523.524 | -0.575 | 0.565 | 0.731 | ns | ns |
| | 0 | 22.5 | -0.828 | 1.814 | 523.178 | -0.456 | 0.648 | 0.789 | ns | ns |
| | | 9 | -1.830 | 0.605 | 523.290 | -3.027 | 0.003 | 0.021 | ** | * |
| | | 13.5 | -0.622 | 1.272 | 523.100 | -0.489 | 0.625 | 0.774 | ns | ns |
| | 10 | 22.5 | -1.056 | 1.465 | 523.065 | -0.721 | 0.471 | 0.679 | ns | ns |
| | | 9 | -1.014 | 0.755 | 523.325 | -1.342 | 0.180 | 0.379 | ns | ns |
| | | 13.5 | NA | NA | NA | NA | NA | NA | ns | ns |
| Abamectin | 60 | 22.5 | -0.784 | 1.276 | 523.117 | -0.615 | 0.539 | 0.721 | ns | ns |
| | | 9 | -0.256 | 0.339 | 1232.728 | -0.754 | 0.451 | 0.665 | ns | ns |
| | | 13.5 | -0.616 | 0.317 | 1227.659 | -1.944 | 0.052 | 0.169 | ns | ns |
| | 0 | 22.5 | -0.520 | 0.330 | 1232.842 | -1.577 | 0.115 | 0.296 | ns | ns |
| | | 9 | 0.428 | 0.324 | 1232.272 | 1.319 | 0.188 | 0.389 | ns | ns |
| | | 13.5 | -0.320 | 0.331 | 1231.803 | -0.966 | 0.334 | 0.573 | ns | ns |
| | 10 | 22.5 | 0.145 | 0.379 | 1232.751 | 0.384 | 0.701 | 0.827 | ns | ns |
| | | 9 | 0.979 | 0.317 | 1232.948 | 3.090 | 0.002 | 0.018 | ** | * |
| | | 13.5 | 0.895 | 0.335 | 1231.342 | 2.673 | 0.008 | 0.044 | ** | * |
| Acifluorfen | 60 | 22.5 | 0.556 | 0.362 | 1206.873 | 1.535 | 0.125 | 0.313 | ns | ns |
| | | 9 | -1.428 | 0.252 | Inf | NA | 0.000 | 0.000 | **** | **** |
| | | 13.5 | -1.976 | 0.233 | Inf | NA | 0.000 | 0.000 | **** | **** |
| | 0 | 22.5 | -2.115 | 0.228 | Inf | NA | 0.000 | 0.000 | **** | **** |
| | | 9 | -0.379 | 0.116 | Inf | NA | 0.001 | 0.012 | ** | * |
| | | 13.5 | -0.071 | 0.128 | Inf | NA | 0.577 | 0.740 | ns | ns |
| | 10 | 22.5 | -0.302 | 0.118 | Inf | NA | 0.011 | 0.058 | * | ns |
| | | 9 | -1.062 | 0.285 | Inf | NA | 0.000 | 0.003 | *** | ** |
| | | 13.5 | -0.789 | 0.287 | Inf | NA | 0.006 | 0.041 | ** | * |
| ha-Cypermeth | 60 | 22.5 | -1.085 | 0.280 | Inf | NA | 0.000 | 0.002 | *** | ** |
| | | 9 | 0.424 | 0.518 | 761.112 | 0.819 | 0.413 | 0.642 | ns | ns |
| | | 13.5 | 0.472 | 0.489 | 760.897 | 0.965 | 0.335 | 0.573 | ns | ns |
| | 0 | 22.5 | 0.304 | 0.579 | 761.984 | 0.526 | 0.599 | 0.748 | ns | ns |
| | | 9 | 0.230 | 0.422 | 761.334 | 0.545 | 0.586 | 0.742 | ns | ns |
| | | 13.5 | 0.639 | 0.372 | 760.458 | 1.719 | 0.086 | 0.245 | ns | ns |
| | 10 | 22.5 | 0.026 | 0.400 | 761.172 | 0.065 | 0.948 | 0.971 | ns | ns |
| | | 9 | -0.047 | 0.499 | 761.443 | -0.095 | 0.925 | 0.962 | ns | ns |
| | | 13.5 | -0.911 | 0.605 | 758.532 | -1.507 | 0.132 | 0.323 | ns | ns |
| Bendiocarb | 60 | 22.5 | -0.383 | 0.414 | 760.391 | -0.924 | 0.356 | 0.595 | ns | ns |
| | | 9 | 0.276 | 0.447 | 2560.611 | 0.617 | 0.537 | 0.721 | ns | ns |
| | | 13.5 | 0.221 | 0.420 | 2560.602 | 0.527 | 0.598 | 0.748 | ns | ns |
| | 0 | 22.5 | 0.567 | 0.426 | 2560.488 | 1.332 | 0.183 | 0.382 | ns | ns |
| | | 9 | -1.029 | 0.293 | 2560.810 | -3.510 | 0.000 | 0.006 | *** | ** |
| | | 13.5 | -1.091 | 0.290 | 2561.548 | -3.769 | 0.000 | 0.003 | *** | ** |
| | 10 | 22.5 | -1.509 | 0.279 | 2560.142 | -5.416 | 0.000 | 0.000 | **** | **** |
| | | 9 | -0.225 | 0.196 | 2560.780 | -1.149 | 0.251 | 0.469 | ns | ns |
| | | 13.5 | -0.042 | 0.184 | 2557.881 | -0.229 | 0.819 | 0.898 | ns | ns |
| Bifenthrin | 60 | 22.5 | -0.040 | 0.229 | 2538.448 | -0.174 | 0.862 | 0.924 | ns | ns |
| | | 9 | -0.698 | 0.338 | 1549.850 | -2.062 | 0.039 | 0.143 | * | ns |
| | | 13.5 | -0.916 | 0.319 | 1544.773 | -2.874 | 0.004 | 0.030 | ** | * |
| | 0 | 22.5 | -1.048 | 0.349 | 1545.153 | -3.001 | 0.003 | 0.022 | ** | * |
| | | 9 | 0.959 | 0.309 | 1549.812 | 3.101 | 0.002 | 0.018 | ** | * |
| | | 13.5 | -0.039 | 0.315 | 1364.002 | -0.125 | 0.900 | 0.947 | ns | ns |
| | 10 | 22.5 | 0.229 | 0.323 | 1549.668 | 0.709 | 0.479 | 0.679 | ns | ns |
| | | 9 | 0.044 | 0.342 | 1494.310 | 0.129 | 0.897 | 0.947 | ns | ns |
| | | 13.5 | 0.592 | 0.406 | 1316.462 | 1.458 | 0.145 | 0.341 | ns | ns |
| Broflanilide | 60 | 22.5 | -0.378 | 0.341 | 1504.897 | -1.107 | 0.268 | 0.486 | ns | ns |
| | | 9 | 0.091 | 1.188 | 40.517 | 0.076 | 0.939 | 0.971 | ns | ns |
| | | 13.5 | -0.345 | 0.883 | 41.662 | -0.391 | 0.698 | 0.827 | ns | ns |
| | 0 | 22.5 | -0.204 | 0.989 | 41.347 | -0.206 | 0.838 | 0.907 | ns | ns |
| | | 9 | 0.649 | 1.393 | 41.312 | 0.466 | 0.644 | 0.787 | ns | ns |
| | | 13.5 | 0.420 | 1.560 | 40.299 | 0.269 | 0.789 | 0.885 | ns | ns |
| | 10 | 22.5 | -0.063 | 1.476 | 40.209 | -0.042 | 0.966 | 0.980 | ns | ns |
| | | 9 | -1.093 | 1.887 | 40.552 | -0.579 | 0.566 | 0.731 | ns | ns |
| | | 13.5 | -0.120 | 1.913 | 40.342 | -0.063 | 0.950 | 0.971 | ns | ns |
| Chlorfenapyr | 60 | 22.5 | -1.583 | 1.968 | 40.209 | -0.804 | 0.426 | 0.649 | ns | ns |
| | | 9 | -0.566 | 0.406 | 1557.429 | -1.392 | 0.164 | 0.361 | ns | ns |
| | | 13.5 | -0.310 | 0.439 | 1553.219 | -0.707 | 0.480 | 0.679 | ns | ns |
| | 0 | 22.5 | -0.825 | 0.375 | 1557.499 | -2.200 | 0.028 | 0.114 | * | ns |
| | | 9 | -0.606 | 0.285 | 1375.366 | -2.124 | 0.034 | 0.129 | * | ns |
| | | 13.5 | -1.111 | 0.283 | 1449.417 | -3.931 | 0.000 | 0.002 | **** | ** |
| | 10 | 22.5 | -0.481 | 0.288 | 1550.113 | -1.673 | 0.095 | 0.260 | ns | ns |
| | | 9 | -0.327 | 0.525 | 1557.370 | -0.623 | 0.534 | 0.721 | ns | ns |
| | | 13.5 | -1.066 | 0.331 | 1470.101 | -3.217 | 0.001 | 0.013 | ** | * |
| dinafop-propa | 60 | 22.5 | -0.761 | 0.343 | 1474.151 | -2.218 | 0.027 | 0.111 | * | ns |
| | | 9 | -0.445 | 0.190 | Inf | NA | 0.019 | 0.088 | * | ns |
| | | 13.5 | -0.966 | 0.182 | Inf | NA | 0.000 | 0.000 | **** | **** |
| | 0 | 22.5 | -1.081 | 0.188 | Inf | NA | 0.000 | 0.000 | **** | **** |
| | | 9 | -0.298 | 0.157 | Inf | NA | 0.058 | 0.182 | ns | ns |
| | | 13.5 | -0.264 | 0.161 | Inf | NA | 0.102 | 0.274 | ns | ns |
| | 10 | 22.5 | -0.334 | 0.175 | Inf | NA | 0.056 | 0.177 | ns | ns |
| | | 9 | -0.603 | 0.224 | Inf | NA | 0.007 | 0.043 | ** | * |
| | | 13.5 | -0.414 | 0.230 | Inf | NA | 0.072 | 0.216 | ns | ns |
| Clothianidin | 60 | 22.5 | -0.701 | 0.238 | Inf | NA | 0.003 | 0.025 | ** | * |
| | | 9 | -0.452 | 0.324 | 1809.004 | -1.393 | 0.164 | 0.361 | ns | ns |
| | | 13.5 | -0.441 | 0.329 | 1814.030 | -1.342 | 0.180 | 0.379 | ns | ns |
| | 0 | 22.5 | -0.817 | 0.343 | 1799.408 | -2.381 | 0.017 | 0.083 | * | ns |
| | | 9 | -0.465 | 0.232 | 1814.034 | -2.004 | 0.045 | 0.158 | * | ns |
| | | 13.5 | -0.693 | 0.247 | 1813.357 | -2.808 | 0.005 | 0.035 | ** | * |
| | 10 | 22.5 | -0.933 | 0.257 | 1772.947 | -3.627 | 0.000 | 0.004 | *** | ** |
| | | 9 | -0.484 | 0.382 | 1814.810 | -1.269 | 0.204 | 0.409 | ns | ns |
| | | 13.5 | -0.820 | 0.400 | 1814.992 | -2.050 | 0.041 | 0.143 | * | ns |
| Cyflumetofen | 60 | 22.5 | -0.218 | 0.363 | 1814.597 | -0.601 | 0.548 | 0.725 | ns | ns |
| | | 9 | 0.643 | 0.370 | 1667.581 | 1.737 | 0.083 | 0.241 | ns | ns |
| | | 13.5 | -0.207 | 0.309 | 1668.719 | -0.670 | 0.503 | 0.702 | ns | ns |
| | 0 | 22.5 | 0.157 | 0.339 | 1667.735 | 0.462 | 0.644 | 0.787 | ns | ns |
| | | 9 | 0.201 | 0.269 | 1667.664 | 0.749 | 0.454 | 0.666 | ns | ns |
| | | 13.5 | 0.282 | 0.255 | 1668.296 | 1.106 | 0.269 | 0.486 | ns | ns |
| | 10 | 22.5 | -0.060 | 0.261 | 1667.869 | -0.230 | 0.818 | 0.898 | ns | ns |
| | | 9 | -1.186 | 0.393 | 1658.351 | -3.016 | 0.003 | 0.021 | ** | * |
| | | 13.5 | -1.209 | 0.385 | 1668.337 | -3.142 | 0.002 | 0.016 | ** | * |
| Deltamethrin | 60 | 22.5 | -1.479 | 0.405 | 1659.075 | -3.657 | 0.000 | 0.004 | *** | ** |
| | | 9 | -0.855 | 0.301 | 826.032 | -2.841 | 0.005 | 0.033 | ** | * |
| | | 13.5 | -0.834 | 0.321 | 826.638 | -2.596 | 0.010 | 0.054 | ** | ns |
| | 0 | 22.5 | -0.966 | 0.333 | 827.811 | -2.904 | 0.004 | 0.028 | ** | * |
| | | 9 | -0.640 | 0.311 | 826.704 | -2.057 | 0.040 | 0.143 | * | ns |
| | | 13.5 | 0.404 | 0.326 | 828.000 | 1.242 | 0.215 | 0.423 | ns | ns |
| | 10 | 22.5 | -1.074 | 0.394 | 827.088 | -2.722 | 0.007 | 0.042 | ** | * |
| | | 9 | -0.341 | 0.517 | 827.849 | -0.659 | 0.510 | 0.708 | ns | ns |
| | | 13.5 | -0.225 | 0.511 | 826.243 | -0.440 | 0.660 | 0.790 | ns | ns |
| Diafenthiuron | 60 | 22.5 | 0.146 | 0.581 | 826.061 | 0.252 | 0.801 | 0.888 | ns | ns |
| | | 9 | -0.427 | 0.362 | 1819.495 | -1.182 | 0.237 | 0.450 | ns | ns |
| | | 13.5 | 0.125 | 0.346 | 1828.992 | 0.361 | 0.718 | 0.842 | ns | ns |
| | 0 | 22.5 | -0.576 | 0.353 | 1828.429 | -1.634 | 0.102 | 0.274 | ns | ns |
| | | 9 | 1.273 | 0.298 | 1791.198 | 4.277 | 0.000 | 0.000 | **** | *** |
| | | 13.5 | 0.199 | 0.271 | 1796.371 | 0.734 | 0.463 | 0.676 | ns | ns |
| | 10 | 22.5 | 0.332 | 0.301 | 1802.014 | 1.103 | 0.270 | 0.486 | ns | ns |
| | | 9 | -0.653 | 0.300 | 1828.631 | -2.175 | 0.030 | 0.119 | * | ns |
| | | 13.5 | -0.243 | 0.312 | 1756.799 | -0.781 | 0.435 | 0.652 | ns | ns |
| Diflubenzuron | 60 | 22.5 | 0.448 | 0.328 | 1702.245 | 1.366 | 0.172 | 0.376 | ns | ns |
| | | 9 | 0.926 | 0.451 | 1596.985 | 2.053 | 0.040 | 0.143 | * | ns |
| | | 13.5 | 0.267 | 0.422 | 1596.447 | 0.634 | 0.526 | 0.721 | ns | ns |
| | 0 | 22.5 | 0.479 | 0.384 | 1597.654 | 1.246 | 0.213 | 0.423 | ns | ns |
| | | 9 | 0.480 | 0.282 | 1581.301 | 1.701 | 0.089 | 0.247 | ns | ns |
| | | 13.5 | 0.055 | 0.290 | 1596.918 | 0.191 | 0.848 | 0.912 | ns | ns |
| | 10 | 22.5 | 0.486 | 0.312 | 1597.890 | 1.557 | 0.120 | 0.305 | ns | ns |
| | | 9 | -0.340 | 0.360 | 1597.528 | -0.944 | 0.345 | 0.588 | ns | ns |
| | | 13.5 | -0.571 | 0.408 | 1596.080 | -1.401 | 0.161 | 0.361 | ns | ns |
| DNOC | 60 | 22.5 | -0.899 | 0.367 | 1597.911 | -2.448 | 0.014 | 0.073 | * | ns |
| | | 9 | 0.650 | 0.577 | 281.797 | 1.126 | 0.261 | 0.483 | ns | ns |
| | | 13.5 | -0.473 | 0.663 | 281.260 | -0.714 | 0.476 | 0.679 | ns | ns |
| | 0 | 22.5 | -0.850 | 0.923 | 281.514 | -0.921 | 0.358 | 0.595 | ns | ns |
| | | 9 | 0.425 | 0.492 | 281.973 | 0.863 | 0.389 | 0.618 | ns | ns |
| | | 13.5 | 0.172 | 0.541 | 281.033 | 0.318 | 0.750 | 0.862 | ns | ns |
| | 10 | 22.5 | -1.875 | 1.780 | 281.662 | -1.053 | 0.293 | 0.517 | ns | ns |
| | | 9 | 2.298 | 0.849 | 281.783 | 2.707 | 0.007 | 0.043 | ** | * |
| | | 13.5 | 0.066 | 1.215 | 280.341 | 0.054 | 0.957 | 0.974 | ns | ns |
| amectin benzo | 60 | 22.5 | -1.166 | 1.336 | 280.797 | -0.872 | 0.384 | 0.618 | ns | ns |
| | | 9 | -0.666 | 0.717 | 224.489 | -0.929 | 0.354 | 0.595 | ns | ns |
| | | 13.5 | -1.183 | 0.650 | 143.536 | -1.820 | 0.071 | 0.216 | ns | ns |
| | 0 | 22.5 | -1.298 | 0.654 | 134.396 | -1.984 | 0.049 | 0.168 | * | ns |
| | | 9 | -0.997 | 0.515 | 346.698 | -1.936 | 0.054 | 0.173 | ns | ns |
| | | 13.5 | -0.206 | 0.634 | 346.554 | -0.325 | 0.746 | 0.862 | ns | ns |
| | 10 | 22.5 | -0.564 | 0.609 | 346.999 | -0.926 | 0.355 | 0.595 | ns | ns |
| | | 9 | -0.673 | 0.942 | 329.475 | -0.714 | 0.476 | 0.679 | ns | ns |
| | | 13.5 | 1.311 | 1.220 | 344.242 | 1.075 | 0.283 | 0.503 | ns | ns |
| Etofenprox | 60 | 22.5 | 0.073 | 1.002 | 346.644 | 0.073 | 0.942 | 0.971 | ns | ns |
| | | 9 | -0.561 | 0.473 | 241.598 | -1.187 | 0.236 | 0.450 | ns | ns |
| | | 13.5 | 0.178 | 0.642 | 242.352 | 0.277 | 0.782 | 0.884 | ns | ns |
| | 0 | 22.5 | 0.062 | 0.497 | 241.658 | 0.125 | 0.901 | 0.947 | ns | ns |
| | | 9 | -0.521 | 0.474 | 242.311 | -1.101 | 0.272 | 0.486 | ns | ns |
| | | 13.5 | 0.164 | 0.533 | 242.749 | 0.308 | 0.758 | 0.867 | ns | ns |
| | 10 | 22.5 | 0.312 | 0.545 | 242.272 | 0.572 | 0.568 | 0.731 | ns | ns |
| | | 9 | 1.607 | 0.717 | 241.217 | 2.241 | 0.026 | 0.111 | * | ns |
| | | 13.5 | 0.377 | 0.892 | 242.498 | 0.422 | 0.673 | 0.802 | ns | ns |
| Fenpyroximate | 60 | 22.5 | 0.450 | 0.731 | 241.717 | 0.615 | 0.673 | 0.721 | ns | ns |
| | | 9 | -0.220 | 0.268 | 984.257 | -0.820 | 0.412 | 0.642 | ns | ns |
| | | 13.5 | -0.353 | 0.276 | 984.708 | -1.281 | 0.200 | 0.407 | ns | ns |
| | 0 | 22.5 | -0.842 | 0.262 | 984.283 | -3.219 | 0.001 | 0.013 | ** | * |
| | | 9 | -0.437 | 0.255 | 985.391 | -1.715 | 0.087 | 0.245 | ns | ns |
| | | 13.5 | -0.115 | 0.330 | 985.915 | -0.350 | 0.727 | 0.848 | ns | ns |
| | 10 | 22.5 | -0.595 | 0.268 | 985.329 | -2.221 | 0.027 | 0.111 | * | ns |
| | | 9 | -0.207 | 0.380 | 984.385 | -0.544 | 0.586 | 0.742 | ns | ns |
| | | 13.5 | -0.008 | 0.376 | 985.148 | -0.020 | 0.984 | 0.989 | ns | ns |
| Flufenoxuron | 60 | 22.5 | -0.136 | 0.422 | 985.854 | -0.322 | 0.747 | 0.862 | ns | ns |
| | | 9 | 0.829 | 0.371 | 368.675 | 2.235 | 0.026 | 0.111 | * | ns |
| | | 13.5 | 0.481 | 0.400 | 368.921 | 1.201 | 0.231 | 0.443 | ns | ns |
| | 0 | 22.5 | 0.622 | 0.372 | 368.312 | 1.670 | 0.096 | 0.261 | ns | ns |
| | | 9 | -0.900 | 0.515 | 362.258 | -1.746 | 0.082 | 0.241 | ns | ns |
| | | 13.5 | -0.547 | 0.445 | 357.860 | -1.230 | 0.219 | 0.424 | ns | ns |
| | 10 | 22.5 | -0.649 | 0.748 | 363.268 | -0.868 | 0.386 | 0.618 | ns | ns |
| | | 9 | 1.057 | 1.074 | 367.761 | 0.984 | 0.326 | 0.564 | ns | ns |
| | | 13.5 | -1.127 | 1.348 | 368.204 | -0.836 | 0.404 | 0.634 | ns | ns |
| ydramethylno | 60 | 22.5 | -1.402 | 1.218 | 368.135 | -1.151 | 0.251 | 0.469 | ns | ns |
| | | 9 | -0.181 | 0.647 | 452.393 | -0.280 | 0.779 | 0.884 | ns | ns |
| | | 13.5 | -0.978 | 0.618 | 452.929 | -1.581 | 0.115 | 0.296 | ns | ns |
| | 0 | 22.5 | -1.390 | 0.598 | 452.987 | -2.322 | 0.021 | 0.092 | * | ns |
| | | 9 | -0.161 | 0.599 | 433.604 | -0.269 | 0.788 | 0.885 | ns | ns |
| | | 13.5 | -0.082 | 0.524 | 421.190 | -0.157 | 0.875 | 0.934 | ns | ns |
| | 10 | 22.5 | 0.449 | 0.569 | 448.707 | 0.789 | 0.431 | 0.649 | ns | ns |
| | | 9 | 0.523 | 0.687 | 419.680 | 0.761 | 0.447 | 0.663 | ns | ns |
| | | 13.5 | -0.874 | 0.710 | 452.951 | -1.232 | 0.219 | 0.424 | ns | ns |
| Imidacloprid | 60 | 22.5 | 0.202 | 0.972 | 452.073 | 0.208 | 0.836 | 0.907 | ns | ns |
| | | 9 | -2.376 | 1.609 | 375.164 | -1.477 | 0.141 | 0.335 | ns | ns |
| | | 13.5 | -2.463 | 1.618 | 373.743 | -1.522 | 0.129 | 0.318 | ns | ns |
| | 0 | 22.5 | NA | NA | NA | NA | NA | NA | ns | ns |
| | | 9 | -1.672 | 1.936 | 359.096 | -0.864 | 0.388 | 0.618 | ns | ns |
| | | 13.5 | -1.564 | 0.985 | 372.622 | -1.588 | 0.113 | 0.296 | ns | ns |
| | 10 | 22.5 | 0.026 | 1.552 | 374.663 | 0.017 | 0.986 | 0.989 | ns | ns |
| | | 9 | -1.356 | 1.044 | 375.969 | -1.299 | 0.195 | 0.401 | ns | ns |
| | | 13.5 | -1.608 | 2.761 | 310.709 | -0.582 | 0.561 | 0.731 | ns | ns |
| Ivermectin | 60 | 22.5 | -0.129 | 1.357 | 374.534 | -0.095 | 0.924 | 0.962 | ns | ns |
| | | 9 | 0.677 | 0.757 | 405.922 | 0.894 | 0.372 | 0.607 | ns | ns |
| | | 13.5 | 0.448 | 0.696 | 405.613 | 0.643 | 0.521 | 0.719 | ns | ns |
| | 0 | 22.5 | 1.016 | 0.719 | 405.995 | 1.414 | 0.158 | 0.359 | ns | ns |
| | | 9 | 0.232 | 0.490 | 404.439 | 0.473 | 0.636 | 0.785 | ns | ns |
| | | 13.5 | -0.669 | 0.466 | 405.013 | -1.436 | 0.152 | 0.351 | ns | ns |
| | 10 | 22.5 | -0.211 | 0.475 | 404.347 | -0.444 | 0.657 | 0.790 | ns | ns |
| | | 9 | -2.767 | 1.075 | 405.734 | -2.575 | 0.010 | 0.057 | * | ns |
| | | 13.5 | -2.205 | 0.980 | 404.094 | -2.251 | 0.025 | 0.110 | * | ns |
| Malaoxon | 60 | 22.5 | -1.343 | 0.998 | 405.240 | -1.346 | 0.179 | 0.379 | ns | ns |
| | | 9 | -0.229 | 0.446 | 903.463 | -0.515 | 0.607 | 0.755 | ns | ns |
| | | 13.5 | -0.467 | 0.453 | 903.506 | -1.030 | 0.303 | 0.529 | ns | ns |
| | 0 | 22.5 | -0.254 | 0.467 | 902.975 | -0.545 | 0.586 | 0.742 | ns | ns |
| | | 9 | -0.839 | 0.353 | 891.599 | -2.375 | 0.018 | 0.083 | * | ns |
| | | 13.5 | -0.710 | 0.363 | 903.475 | -1.955 | 0.051 | 0.169 | ns | ns |
| | 10 | 22.5 | -0.299 | 0.412 | 903.301 | -0.726 | 0.468 | 0.679 | ns | ns |
| | | 9 | -0.387 | 0.486 | 903.676 | -0.796 | 0.426 | 0.649 | ns | ns |
| | | 13.5 | -1.183 | 0.467 | 895.731 | -2.535 | 0.011 | 0.060 | * | ns |
| Novaluron | 60 | 22.5 | -0.780 | 0.528 | 903.760 | -1.477 | 0.140 | 0.335 | ns | ns |
| | | 9 | -1.583 | 0.586 | 1166.937 | -2.703 | 0.007 | 0.043 | ** | * |
| | | 13.5 | -0.634 | 0.615 | 1166.335 | -1.032 | 0.302 | 0.529 | ns | ns |
| | 0 | 22.5 | -1.632 | 0.586 | 1165.517 | -2.783 | 0.005 | 0.037 | ** | * |
| | | 9 | -0.322 | 0.409 | 1165.168 | -0.788 | 0.431 | 0.649 | ns | ns |
| | | 13.5 | 1.019 | 0.470 | 1165.623 | 2.167 | 0.030 | 0.119 | * | ns |
| | 10 | 22.5 | -0.089 | 0.419 | 1165.756 | -0.212 | 0.832 | 0.907 | ns | ns |
| | | 9 | -3.067 | 0.567 | 1166.914 | -5.412 | 0.000 | 0.000 | **** | **** |
| | | 13.5 | -2.426 | 0.547 | 1165.656 | -4.431 | 0.000 | 0.000 | **** | *** |
| Permethrin | 60 | 22.5 | -1.919 | 0.585 | 1165.769 | -3.279 | 0.001 | 0.012 | ** | * |
| | | 9 | 0.004 | 0.290 | 739.636 | 0.013 | 0.989 | 0.989 | ns | ns |
| | | 13.5 | -0.059 | 0.302 | 753.823 | -0.194 | 0.846 | 0.912 | ns | ns |
| | 0 | 22.5 | -0.469 | 0.346 | 737.413 | -1.357 | 0.175 | 0.378 | ns | ns |
| | | 9 | -0.900 | 0.338 | 729.697 | -2.668 | 0.008 | 0.045 | ** | * |
| | | 13.5 | -0.699 | 0.358 | 741.944 | -1.954 | 0.051 | 0.169 | ns | ns |
| | 10 | 22.5 | -1.216 | 0.401 | 754.995 | -3.034 | 0.002 | 0.021 | ** | * |
| | | 9 | 0.487 | 0.535 | 749.474 | 0.910 | 0.363 | 0.597 | ns | ns |
| | | 13.5 | -0.351 | 0.659 | 743.567 | -0.533 | 0.594 | 0.748 | ns | ns |
| eronyl-butoxi | 60 | 22.5 | -0.332 | 0.524 | 754.938 | -0.633 | 0.527 | 0.721 | ns | ns |
| | | 9 | -0.401 | 0.287 | 2183.956 | -1.396 | 0.163 | 0.361 | ns | ns |
| | | 13.5 | -0.922 | 0.275 | 2182.763 | -3.351 | 0.001 | 0.010 | *** | ** |
| | 0 | 22.5 | -0.729 | 0.267 | 2184.000 | -2.734 | 0.006 | 0.041 | ** | * |
| | | 9 | 0.479 | 0.200 | 2183.732 | 2.392 | 0.017 | 0.083 | * | ns |
| | | 13.5 | 0.056 | 0.184 | 2182.909 | 0.304 | 0.761 | 0.867 | ns | ns |
| | 10 | 22.5 | 0.082 | 0.182 | 2182.727 | 0.449 | 0.654 | 0.790 | ns | ns |
| | | 9 | -0.075 | 0.224 | 2183.940 | -0.333 | 0.739 | 0.859 | ns | ns |
| | | 13.5 | -0.131 | 0.226 | 2183.664 | -0.581 | 0.562 | 0.731 | ns | ns |
| rimiphos-meth | 60 | 22.5 | -0.251 | 0.198 | 2183.517 | -1.270 | 0.204 | 0.409 | ns | ns |
| | | 9 | -1.143 | 0.207 | Inf | NA | 0.000 | 0.000 | **** | **** |
| | | 13.5 | -1.102 | 0.203 | Inf | NA | 0.000 | 0.000 | **** | **** |
| | 0 | 22.5 | -1.155 | 0.207 | Inf | NA | 0.000 | 0.000 | **** | **** |
| | | 9 | -0.201 | 0.131 | Inf | NA | 0.126 | 0.313 | ns | ns |
| | | 13.5 | -0.108 | 0.136 | Inf | NA | 0.426 | 0.649 | ns | ns |
| | 10 | 22.5 | -0.313 | 0.125 | Inf | NA | 0.012 | 0.064 | * | ns |
| | | 9 | -0.341 | 0.164 | Inf | NA | 0.037 | 0.139 | * | ns |
| | | 13.5 | -0.360 | 0.167 | Inf | NA | 0.031 | 0.121 | * | ns |
| Pyriproxyfen | 60 | 22.5 | -0.307 | 0.162 | Inf | NA | 0.059 | 0.182 | ns | ns |
| | | 9 | -0.566 | 0.301 | 1578.606 | -1.880 | 0.060 | 0.185 | ns | ns |
| | | 13.5 | -0.675 | 0.312 | 1604.939 | -2.166 | 0.030 | 0.119 | * | ns |
| | 0 | 22.5 | -1.111 | 0.300 | 1537.159 | -3.709 | 0.000 | 0.003 | *** | ** |
| | | 9 | -0.016 | 0.250 | 1617.969 | -0.064 | 0.949 | 0.971 | ns | ns |
| | | 13.5 | -0.057 | 0.220 | 1618.367 | -0.257 | 0.798 | 0.888 | ns | ns |
| | 10 | 22.5 | 0.094 | 0.246 | 1617.523 | 0.381 | 0.703 | 0.827 | ns | ns |
| | | 9 | -0.153 | 0.257 | 1617.093 | -0.598 | 0.550 | 0.725 | ns | ns |
| | | 13.5 | 0.027 | 0.241 | 1611.785 | 0.111 | 0.912 | 0.955 | ns | ns |
| Sherlock | 60 | 22.5 | 0.185 | 0.269 | 1606.719 | 0.689 | 0.491 | 0.690 | ns | ns |
| | | 9 | 0.472 | 0.267 | Inf | NA | 0.077 | 0.231 | ns | ns |
| | | 13.5 | 0.217 | 0.248 | Inf | NA | 0.382 | 0.618 | ns | ns |
| | 0 | 22.5 | 0.304 | 0.248 | Inf | NA | 0.219 | 0.424 | ns | ns |
| | | 9 | -0.753 | 0.233 | Inf | NA | 0.001 | 0.013 | ** | * |
| | | 13.5 | -0.318 | 0.218 | Inf | NA | 0.146 | 0.341 | ns | ns |
| | 10 | 22.5 | -0.849 | 0.232 | Inf | NA | 0.000 | 0.004 | *** | ** |
| | | 9 | -0.826 | 0.391 | Inf | NA | 0.035 | 0.130 | * | ns |
| | | 13.5 | 0.043 | 0.296 | Inf | NA | 0.883 | 0.939 | ns | ns |
| Spinosad | 60 | 22.5 | -1.552 | 0.378 | Inf | NA | 0.000 | 0.001 | **** | *** |
| | | 9 | -0.843 | 0.588 | 353.390 | -1.435 | 0.152 | 0.351 | ns | ns |
| | | 13.5 | -1.286 | 0.537 | 353.262 | -2.393 | 0.017 | 0.083 | * | ns |
| | 0 | 22.5 | -1.427 | 0.572 | 353.048 | -2.496 | 0.013 | 0.067 | * | ns |
| | | 9 | -0.171 | 0.660 | 353.120 | -0.259 | 0.796 | 0.888 | ns | ns |
| | | 13.5 | -1.075 | 0.627 | 353.234 | -1.714 | 0.087 | 0.245 | ns | ns |
| | 10 | 22.5 | -0.531 | 0.624 | 353.196 | -0.851 | 0.395 | 0.625 | ns | ns |
| | | 9 | 0.152 | 0.613 | 354.528 | 0.248 | 0.804 | 0.888 | ns | ns |
| | | 13.5 | 0.025 | 1.087 | 353.051 | 0.023 | 0.982 | 0.989 | ns | ns |
| Teflubenzuron | 60 | 22.5 | -1.046 | 0.771 | 353.873 | -1.357 | 0.176 | 0.378 | ns | ns |
| | | 9 | -0.656 | 0.426 | 545.824 | -1.541 | 0.124 | 0.313 | ns | ns |
| | | 13.5 | -0.332 | 0.483 | 545.280 | -0.687 | 0.492 | 0.690 | ns | ns |
| | 0 | 22.5 | -0.565 | 0.440 | 545.739 | -1.282 | 0.200 | 0.407 | ns | ns |
| | | 9 | 0.354 | 0.465 | 545.347 | 0.761 | 0.447 | 0.663 | ns | ns |
| | | 13.5 | 0.645 | 0.575 | 544.597 | 1.122 | 0.262 | 0.483 | ns | ns |
| | 10 | 22.5 | 0.346 | 0.578 | 545.446 | 0.599 | 0.549 | 0.725 | ns | ns |
| | | 9 | 0.801 | 0.535 | 545.538 | 1.497 | 0.135 | 0.327 | ns | ns |
| | | 13.5 | 1.177 | 0.742 | 545.174 | 1.587 | 0.113 | 0.296 | ns | ns |
| Tralopyril | 60 | 22.5 | 0.548 | 0.690 | 545.892 | 0.794 | 0.427 | 0.649 | ns | ns |
| | | 9 | -0.393 | 0.351 | 1676.960 | -1.119 | 0.263 | 0.483 | ns | ns |
| | | 13.5 | -0.616 | 0.317 | 1675.190 | -1.945 | 0.052 | 0.169 | ns | ns |
| | 0 | 22.5 | -0.604 | 0.307 | 1676.031 | -1.970 | 0.049 | 0.168 | * | ns |
| | | 9 | 1.645 | 0.389 | 1676.354 | 4.227 | 0.000 | 0.001 | **** | *** |
| | | 13.5 | 0.405 | 0.235 | 1676.586 | 1.726 | 0.085 | 0.244 | ns | ns |
| | 10 | 22.5 | -0.173 | 0.189 | 1676.079 | -0.916 | 0.360 | 0.595 | ns | ns |
| | | 9 | 0.908 | 0.389 | 1676.167 | 2.335 | 0.020 | 0.089 | * | ns |
| | | 13.5 | 0.469 | 0.332 | 1675.290 | 1.412 | 0.158 | 0.359 | ns | ns |
| Transfluthrin | 60 | 22.5 | 0.221 | 0.360 | 1620.006 | 0.615 | 0.538 | 0.721 | ns | ns |
